# VDAC1 selective molecules promote patients’-derived cancer organoids death through mitochondrial-dependent metabolic interference

**DOI:** 10.1101/2023.12.04.569205

**Authors:** Stefano Conti Nibali, Silvia De Siervi, Enrico Luchinat, Andrea Magrì, Lorenza Brocca, Stefania Mantovani, Barbara Oliviero, Mario U. Mondelli, Vito De Pinto, Cristian Turato, Cristina Arrigoni, Marco Lolicato

## Abstract

In the continuous pursuit of advanced cancer therapeutics, our research unveils the potential to selectively target Voltage-Dependent Anion-selective Channel isoform 1 (VDAC1), a pivotal component in cellular metabolism and apoptosis. VDAC1’s role in metabolic rewiring and its subsequent prominence in many cancer types offer a unique intervention point. The incorporation of a systematic, *in silico* to *in vitro* methodology identified novel VA (**<u>V</u>**DAC-**<u>A</u>**ntagonist) molecules with the capability to selectively bind to VDAC1, displaying a substantial specificity towards cancer cells while sparing healthy ones.

This research first led to the revelation of a specialized VDAC1 pocket, which accommodates the binding of these VA molecules, thereby instigating a selective displacement of NADH. The coenzyme is a critical metabolic substrate, and its displacement ensues in notable mitochondrial distress and a reduction in cell proliferation, specifically in cancer cells. Furthermore, meticulous analysis using organoids derived from intrahepatic cholangiocarcinoma patients (iCCA) demonstrated a dose-dependent reduction in cell viability upon treatment with VA molecules, correlating with the findings from commercial cell lines.

Interestingly, VA molecules significantly reduced cell viability and demonstrated a lower impact on healthy cells than conventional treatments like gemcitabine. This differential impact is possibly due to the elevated expression of VDAC1 in various cancer cell lines, rendering them more susceptible to metabolic disruptions induced by VA molecules.

This endeavor uncovers a multifaceted approach to cancer treatment, involving meticulous targeting of metabolic gatekeepers like VDAC1 using novel entities, thereby paving the way for developing more selective and refined cancer therapeutics. The identified VA molecules, albeit in the nascent stages, represent promising candidates for further optimization and development, potentially revolutionizing treatment modalities in cancer therapy through precise metabolic interventions.

## Introduction

Cancer, a complex and devastating disease, continues to challenge medical researchers worldwide. Successful development of cancer therapy has roots in understanding derailed physiological mechanisms observed in cancer cells, including metabolic rewiring, used by tumor cells to escape immunosurveillance thus leading to apoptosis evasion and uncontrolled proliferation (Dang and Kim, 2018).

The Warburg Effect, a metabolic alteration observed in many cancer cell types, has garnered significant attention as a possible cancer cell-selective target mechanism. In this metabolic alteration, glucose metabolism is predominantly directed toward biosynthetic pathways required for rapid cell proliferation and growth. In this context, both glycolysis and oxidative phosphorylation are simultaneously enhanced to thrive cancer cell proliferation (DeBerardinis and Chandel, 2016) by tampering with mitochondrial metabolism.

The maintenance of mitochondrial metabolism relies on the balanced flow of respiratory substrates, nucleotides (ATP, ADP), and ions into and out of mitochondria. Voltage-Dependent Anion-selective Channels (VDAC) in the outer mitochondrial membrane (OMM) play a crucial role in maintaining cell bioenergetic, transporting most anionic metabolites and small ions in and out the mitochondria. In mammals, three different isoforms (VDAC1, VDAC2 and VDAC3) exist and are characterized by a molecular weight of 28–32 kDa and by about 70% sequence similarity (Messina et al., 2012). Among the isoforms, VDAC1 is the regulation of mitochondria-mediated apoptosis. Specifically, VDAC1 partners with anti- and pro-apoptotic proteins (Camara et al., 2017). Under physiological conditions, VDAC1 interacts with anti-apoptotic Bcl-2 family members to inhibit apoptosis, while in the presence of cellular stress or apoptotic stimuli the channel drives the activation of the intrinsic apoptotic pathway and eventually leads to programmed cell death.

There is mounting evidence that VDAC1 interacts with the N-terminal portion of the glycolytic enzymes Hexokinases (HKs) (Haloi et al., 2021a; Magrì et al., 2021). The interaction between HKs and VDAC1 has a dual effect: (1) it prevents the interaction of the porin with pro-apoptotic factors (Vyssokikh and Brdiczka, 2003) and, (2) provides tumor cells with a metabolic advantage through the “Warburg effect”. Colocalization of HK proteins and VDAC1 gives HKs preferential access to the ATP produced within mitochondria. Interestingly, high expression of both VDAC1 and HKs has been observed in many cancer types (Fang et al., 2022; Gao et al., 2015; Haloi et al., 2021b; Magri et al., 2021; Pittala et al., 2018; Wang et al., 2016; Zhang et al., 2016). This allows cancer cells to maintain a high rate of glycolysis, even under aerobic conditions, which provides a continuous supply of building blocks for rapid cell proliferation (Pedersen, 2008). Overall, the altered VDAC1 expression and function appear related to highly metabolically active and energy-demanding cancer cells, suggesting that the channel may act as an oncogene prompting the tumor progression by different molecular mechanisms, including the deregulation of apoptosis (Mazure, 2017; Shoshan-Barmatz et al., 2015; Shoshan-Barmatz and Golan, 2012). Thus, the specific role of VDAC1 in this pathological context together with its shown druggability strengthened the suitability of these proteins as pharmacological targets (Magri et al., 2018; Reina and De Pinto, 2017). To this date several molecules and peptides have been developed to modulate VDAC1 activity to impair cancer cells’ energy homeostasis, proliferation and apoptosis. To date, three distinct class of molecules or peptides have been tested aimed at i) interfere with the VDAC1-HKs interaction; ii) modulate the electrophysiological activity of the channel; iii) affect the expression levels of VDAC1 gene (Magri et al., 2018). However, in most cases, the identified molecules have not met the need of a cancer cell-specific action and/or their binding pocket has not been fully characterized.

In light of this, through an initial *in silico* approach and subsequent thorough biochemical binding assays and NMR spectroscopy, we have identified a set of novel chemical entities, hereafter identified *VA, i.e.* **<u>V</u>**DAC-**<u>A</u>**ntagonist, sharing a three-ring architecture able to specifically bind VDAC1 in a defined pocket shared, but not identical, to that one of NADH. Binding competition assays revealed that VA molecules compete with NADH in a dose-response manner, resulting in the interference of its binding to the channel.

Also, cell proliferation and promotion of cell death in living organoid settings resulted reduced depending on the applied dose of VA molecules. Furthermore, the displacement of NADH from VDAC1 results in lower mitochondrial oxygen consumption, suggesting mitochondrial distress. Altogether these findings suggest that cancer cell proliferation can be effectively halted by VA molecules treatment, leaving the surrounding healthy cells unaffected. This result strengthens the observations about the important role of VDAC as governor of cell fate.

## Results

### High-throughput screening identifies five VDAC1 binders

The overexpression of VDAC1 in highly glycolytic cancer cells prompted us to develop a workflow for the identification of novel chemical entities able to decrease energy production in cancer cells. VDAC1 is a β−barrel channel constituted by 19 anti-parallel β-strands and a N-terminal α-helix (Fig. 1A). The N-terminal helix is a highly mobile element of the protein fundamental for the channel gating (Najbauer et al., 2022) and for the interaction of VDAC1 with the cytosolic protein partners (Manzo et al., 2018). To date, the N-terminal helix has been structurally observed inside the β-barrel where it separates the channel lumen from a small cavity towards the barrel. Part of this groove accommodates the nicotinamide moiety of NADH (Fig. 1B), the main oxidizing power accumulator from the TCA cycle inside the mitochondria.

**Figure 1.**
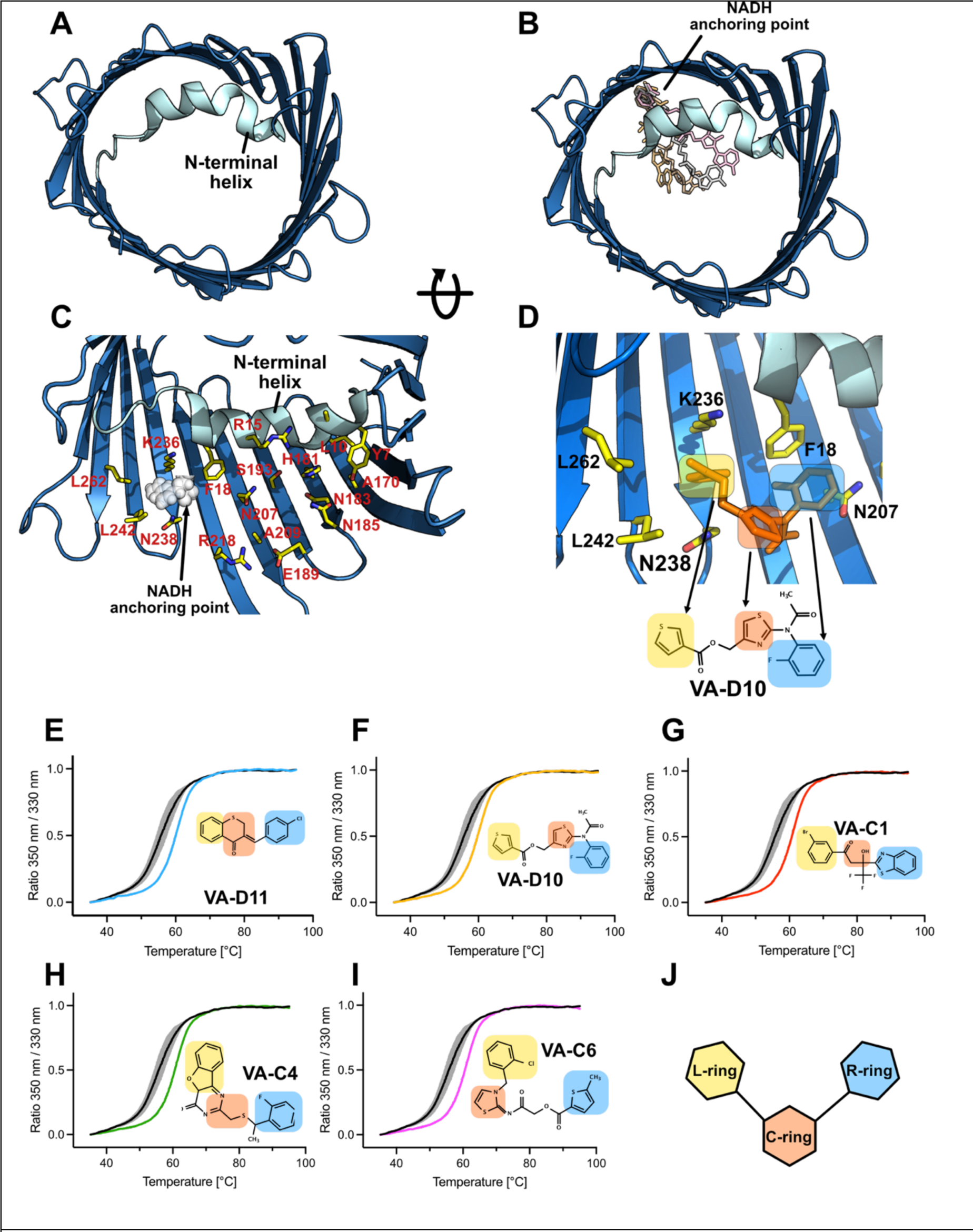
VDAC1 architecture and small molecule identification. (A) Cartoon representation of mouse VDAC1 crystal structure (PDB: 3EMN). In blue is identified the β-strand topology of the channel, and in cyan, its N-terminal α-helix. (B) Selected poses of NADH bound to the channel (PDB: 6TIR), with its nicotinamide anchoring point highlighted. (C) The large groove between VDAC1 N-terminal helix and the β-barrel used as a target for the AtomNet drug discovery campaign. NADH anchoring point is indicated. (D) Close-up view of the putative VA compounds binding pocket, showing the VA-D10 molecule docked in the pocket and assuming a V-shaped conformation. (E-I) Representative denaturation curves of mVDAC1 (0.9 mg/ml) in LDAO after treatment with different VA molecules at a final concentration of 1 mM. Black curves represent the denaturation profile of the protein pre-treated with DMF (0.5 %), and the colored ones the protein pre-treated with the indicated VA molecules. Temperature denaturation inflection point (Ti) shifts towards higher temperatures in the presence of VA molecules VA-D11 (+ 4.66 ± 0.11); VA-D10 (+ 4.93 ± 0.09); VA-C1 (+ 4.58 ± 0.15); VA-C4 (+ 3.58 ± 0.16); VA-C6 (+ 4.59 ± 0.10), indicating protein stabilization/binding. Data are expressed as mean ± SEM (n = 3).

The rationale behind the drug discovery was to hamper the interaction of VDAC1 N-terminal helix with anti-apoptotic factors by stabilizing its interaction within the barrel cavity. Thus, we partnered with Atomwise Inc. (San Francisco, CA) for the *in silico* search of molecules able to staple VDAC1 N-terminal helix within the cavity of the β-barrel. We identified a putative large binding area composed of the N-terminal helix (residues 1-21) and β-strands 12-18 (Fig. 1A, C). This region includes the NADH anchoring point (Fig. 1B, C) (Bohm et al., 2020).

Atomwise technology screened a proprietary chemical library of millions of compounds and identified 70 hits that were experimentally tested for *in vitro* binding with a suite of assays.

First, in order to maximize the throughput, we tested the seventy molecules by measuring the VDAC1 protein thermal denaturation in absence and in presence of the compounds, relying on the correlation between an increase of the apparent denaturation temperature of the purified VDAC1 protein and a stabilization effect given by the bound molecule. Out of the seventy molecules, five compounds (hereafter called VA, **<u>V</u>**DAC-**<u>A</u>**ntagonist) stabilized the apparent VDAC1 unfolding temperature by >3°C (Fig. 1E-I), a significant change consistent with protein-molecule binding(Huynh and Partch, 2015).

The five selected molecules were graduated to a biophysical characterization to confirm *in vitro* binding to the purified VDAC1.

The chemical identity of the five molecules was confirmed by ^1^H-NMR (Suppl. Fig. S1). ^19^F NMR was additionally recorded for the molecules VA-D10, VA-C4 and VA-C1 which present three (VA-C1) and one (VA-D10 and VA-C4) fluorine atom, respectively.

The ^19^F nucleus is highly sensitive to changes in the chemical environment and is absent in biological macromolecules and detergents, making it an ideal probe for monitoring protein-ligand interactions in complex mixtures (Buchholz and Pomerantz, 2021; Dalvit and Vulpetti, 2019). Therefore, the presence of fluorine atoms was exploited to test the direct binding to VDAC1 by monitoring the ^19^F signal of each compound in a micelle-containing buffer in the absence and in the presence of VDAC1. All three compounds experienced line broadening even without VDAC1, likely due to non-specific interaction with the micelles. The presence of VDAC1 caused further line broadening (VA-C1 and VA-D10) and/or shifting (VA-C4 and VA-D10) of the ^19^F resonance in the spectrum of each molecule (Fig. 2), thereby indicating direct binding to the protein.

**Figure 2.**
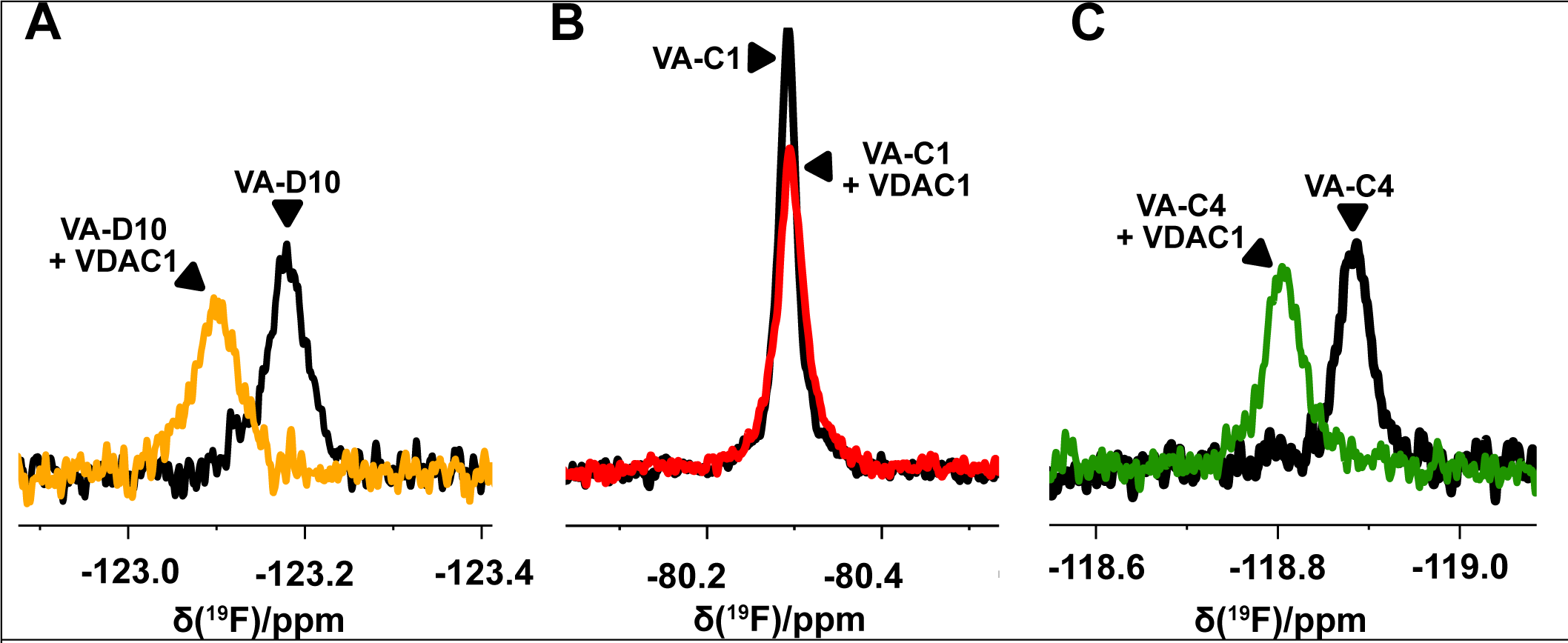
VA molecule binding measured by ^19^F NMR. (A-C) ^19^F NMR spectra of indicated VA molecules in LDAO buffer (black lines) and in the presence of 5μM mVDAC1 (colored lines). VA-D10 (A) and VA-C4 (C) significantly move their chemical shifts upon VDAC1 binding. VA-C1 (B) shows a reduction and broadening of its peak upon VDAC1 interaction.

These experiments confirmed that out of seventy molecules obtained from the *in silico* screening five are *in vitro* binders, so we focused on this set for biophysical characterization of the binding mode and affinity.

### VA molecules share part of the binding site with NADH and bind VDAC1 with micromolar affinity

We docked the VA molecules into the target binding region identified in the *in silico* screening. All five molecules share a “*V shape”* conformation and three aromatic rings in their structure (Fig. 1D-J). For simplicity we call the three rings L-(left), C-(central) and R-(right). All five molecules present the three-ringed structure, except for VA-C1 which has a pseudo-C-ring conformation.

From docking we obtained several poses revealing that all the molecules shared part of the NADH binding pocket (Fig. 1D). In particular, VA molecules are framed by residues F18 (π−π interaction with R-ring), N207, R218 (cation-π interaction with C-ring), K236 (cation-π interaction with L-ring) and N238. Considering the dynamics of the NMR structure, the NADH nicotinamide moiety seems to overlap with only the L-ring of the docked VA molecules and only marginally involves the N-terminal helix, which makes more contacts with the more flexible part of NADH (Bohm et al., 2020). In contrast, the three-ringed architecture of the VA molecules enables a firm anchoring of the N-terminal helix to the barrel by the insertion of the helix F18 into the V shaped compound (Fig. 1D). To test the hypothesis that VA molecules and NADH share the binding pocket, we carry out a Fluorescence Polarization (FP) assay based on the intrinsic fluorescence of NADH molecules (λ_ex_=355, λ_em_=460 nm). We measured the change in fluorescence polarization upon binding of NADH to purified VDAC1 and we fitted the data with a Hill curve obtaining the apparent affinity (K_D_=6.8±0.06, Fig. 3A). We then used a concentration of NADH of 5μM to perform a competition assay in the presence of increasing VA molecules concentrations (Fig. 3B-F). All five molecules showed a decrease in the fluorescence polarization signal in a dose-response manner, confirming the competition with NADH binding with an apparent K_D_ in the micromolar range (K_D_VA-D11=15.9±0.7, K_D_VA-D10=18.5±0.2, K_D_VA-C1=18.7±0.3, K_D_VA-C4=19.4±0.4, K_D_VA-C6=18.7±0.6).

**Figure 3.**
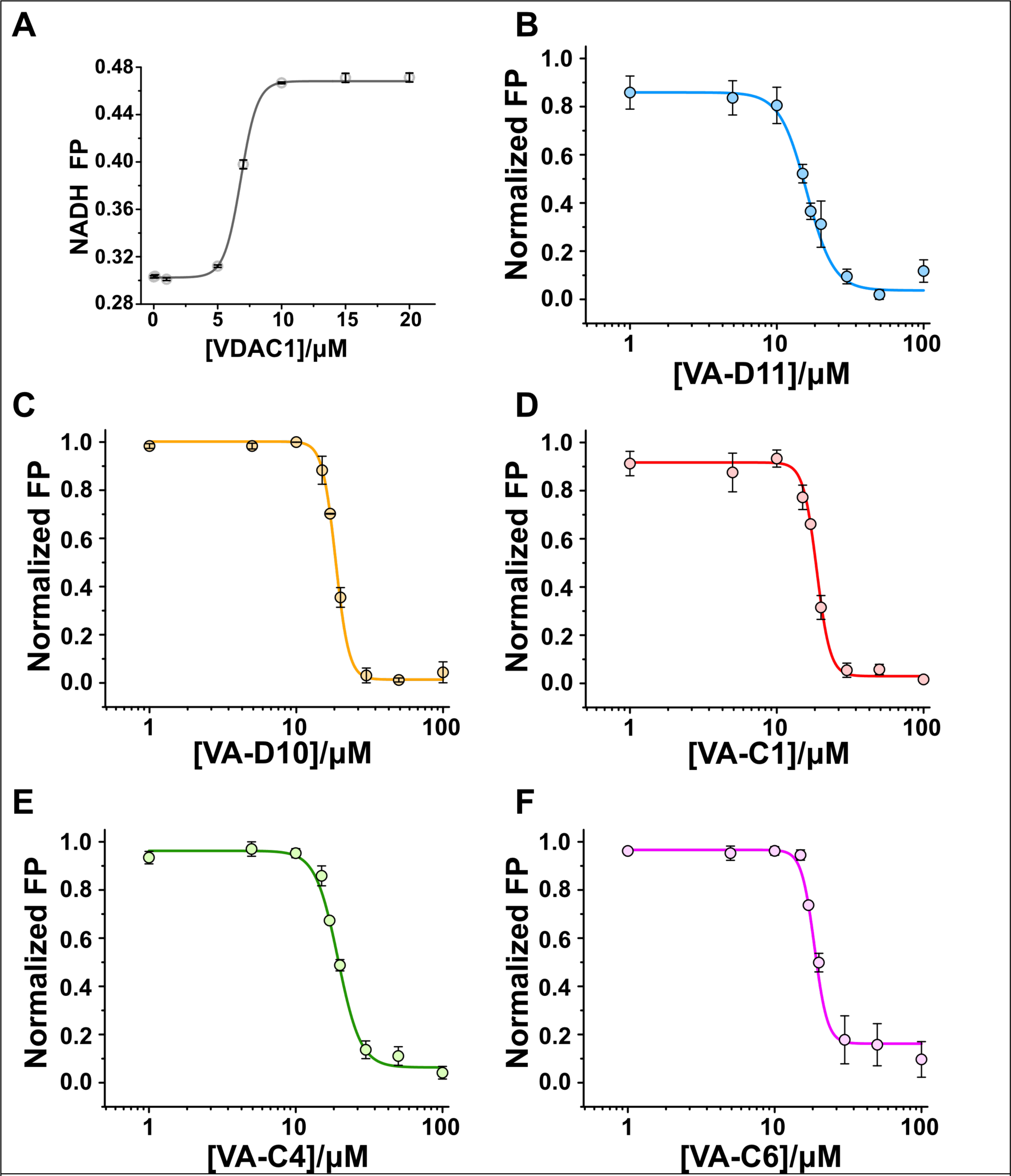
Direct binding of VA molecules to VDAC1 revealed by Fluorescence Polarization competition assay. (A) Binding of the NADH (at 5 µM) to increasing concentrations of mVDAC1 measured by fluorescence polarization (FP). 7 µM is the protein concentration used in all experiments. (B-F) Displacement curves of the NADH (5 µM) from mVDAC1 with increased concentrations of VA-molecules. All the five molecules are able to compete with NADH binding with an apparent KD in the micromolar range (KDVA-D11=15.9±0.64, KDVA-D10=18.5±0.21, KDVA-C1=18.7±0.28, KDVA-C4=19.4±0.36, KDVA-C6=18.7±0.51). Data are expressed as mean ± SEM (n = 3).

Based on the key residues identified in the docking simulations we designed mutations in VDAC1 to confirm the binding and test the importance of the putative interactions.

We tested mutations that supposedly either stabilize or destabilize the binding of VA molecules by means of thermal denaturation assays (Fig. 4A-F) on purified VDAC1. If the molecules bind in the V-shape conformation, introducing positive charges or aromatic residues in place of neutral amino acids (N207R, N238K/F) should strengthen the interactions with the aromatic rings. On the opposite, removing the key π−π interaction with R-ring (F18A) and the cation-π interaction with L-ring (K236A), should lower the affinity of the molecule. To quantify the effect of the mutations on the binding we calculated the difference of the inflection temperature (ΔΔTi) of each mutant in absence or presence of the molecules (Δτi_mut(VA_mol)_-ΔTi_mut(DMSO)_) compared to those measured for VDAC1 wild-type (ΔTi_wt(VA_mol)_-ΔTi_wt(DMSO)_). A positive value was observed for mutants N238K, N238R, N207R, N238F, R218Y consistent with a higher thermostability of these VDAC1 mutants when bound to the molecules (Fig 4). On a opposite trend, alanine substitutions of charged and aromatic residues (K236A, F18A) resulted in a negative difference of the relative unfolding temperatures for all the five molecules. The significance of the differences measured was also confirmed by the unchanged Ti of the mutant in control conditions compared to the wild-type (Table S1), suggesting that these mutations *per se* do not alter the overall stability of the protein.

**Figure 4.**
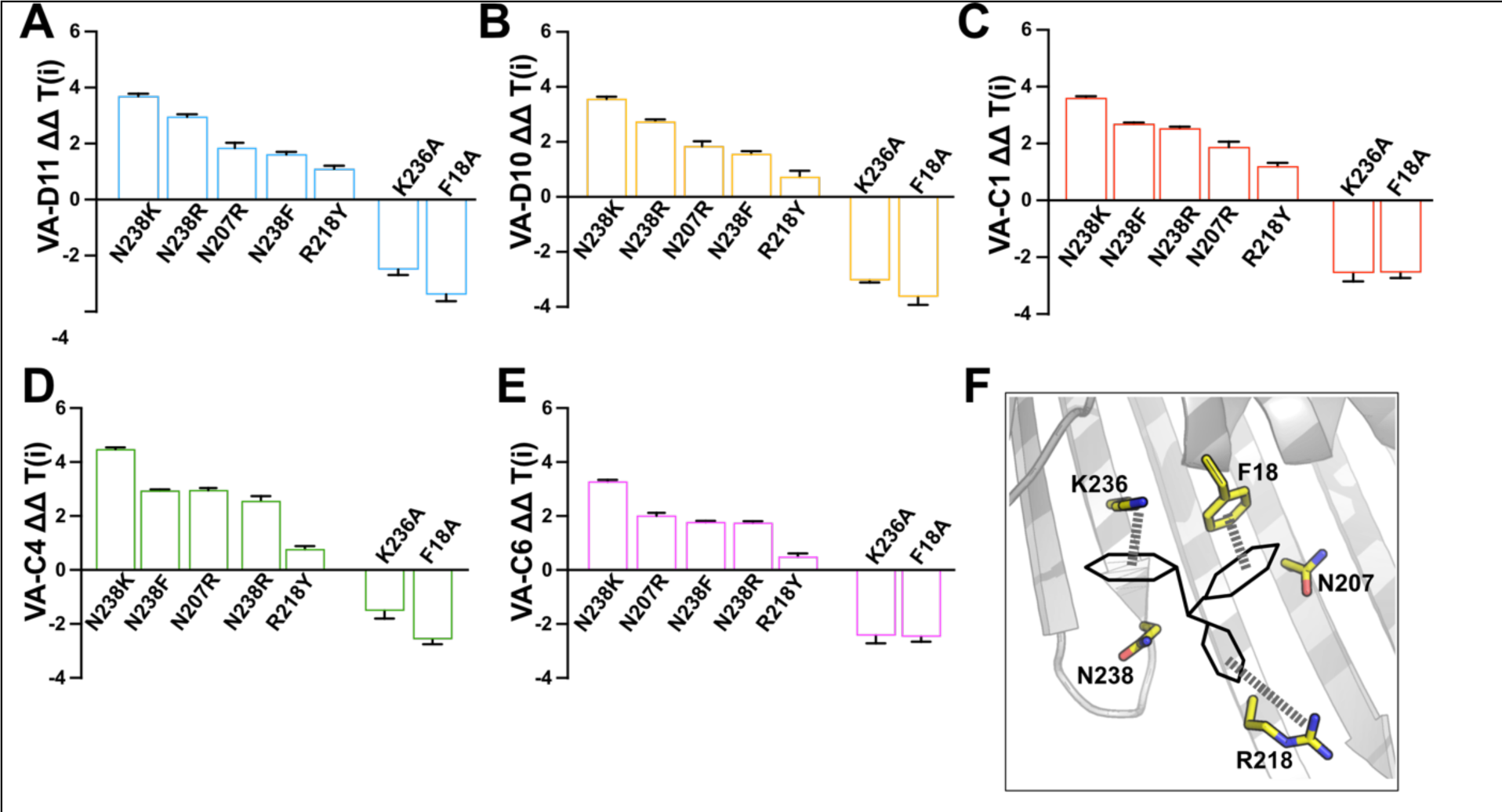
Identification of VA molecules framing residues. (A-E). ΔΔ inflection temperatures (ΔΔTi) values were obtained by subtracting the ΔTi of mVDAC1 mutants from the ΔTi of wild-type protein. Positive or negative ΔΔTi values show the effect of stabilizing and destabilizing mutations on inflection temperature after treatment with VA molecules, respectively. Data are expressed as mean ± SEM (n = 3). (F) Schematic representation of the V-shaped VA compounds and the interactions between VDAC1 residues and the three aromatic rings of the molecules. The thermal denaturation assays confirmed that a cation-π interaction between K236 and VA-L rings and a π−π stack between F18 and VA-R rings are essential for binding. The binding can be further improved by introducing additional π interactions in the neighboring residues.

The thermal denaturation assays confirmed the docking generated hypothesis of the binding mode of the VA molecules showing a decreased apparent affinity for the destabilizing mutations and an improved binding for the stabilizing mutations (Table 1).

**Table 1.** Inflection temperature (T(i)) values. The table shows the T(i) values of the recombinant proteins mVDAC1 wilt-type (WT) and mutants treated with the different small molecules at the final concentration of 1 mM in DMF. Data are reported as mean ± SEM of n=3 independent experiments.

| *T(i) [°C]* | *WT* | *N238K* | *N238F* | *N238R* | *R218Y* | *N207R* | *K236A* | *F18A* |
| --- | --- | --- | --- | --- | --- | --- | --- | --- |
| **VA-D11** | 60.2 ± 0.1 | 64.5 ± 0.1 | 63.1 ± 0.1 | 63.1 ± 0.1 | 62.3 ± 0.1 | 62.9 ± 0.1 | 57.9 ± 0.1 | 57.2 ± 0.1 |
| **VA-D10** | 60.5 ± 0.1 | 64.4 ± 0.1 | 63.3 ± 0.1 | 63.1 ± 0.1 | 62.2 ± 0.2 | 62.9 ± 0.1 | 57.9 ± 0.1 | 57.3 ± 0.1 |
| **VA-C1** | 60.2 ± 0.1 | 64.0 ± 0.1 | 64.0 ± 0.1 | 62.5 ± 0.1 | 62.3 ± 0.1 | 63.2 ± 0.1 | 57.3 ± 0.1 | 58.0 ± 0.1 |
| **VA-C4** | 59.2 ± 0.1 | 64.0 ± 0.1 | 63.3 ± 0.1 | 61.6 ± 0.1 | 60.8 ± 0.1 | 63.0 ± 0.1 | 57.1 ± 0.1 | 57.9 ± 0.1 |
| **VA-C6** | 60.2 ± 0.1 | 63.8 ± 0.1 | 63.2 ± 0.1 | 61.8 ± 0.1 | 61.6 ± 0.1 | 63.0 ± 0.1 | 57.0 ± 0.1 | 58.1 ± 0.1 |

To quantify these mutations’ effect more rigorously, we performed FP assays of the two VDAC1 mutants, K236A and N238K (Fig. 5).

**Figure 5.**
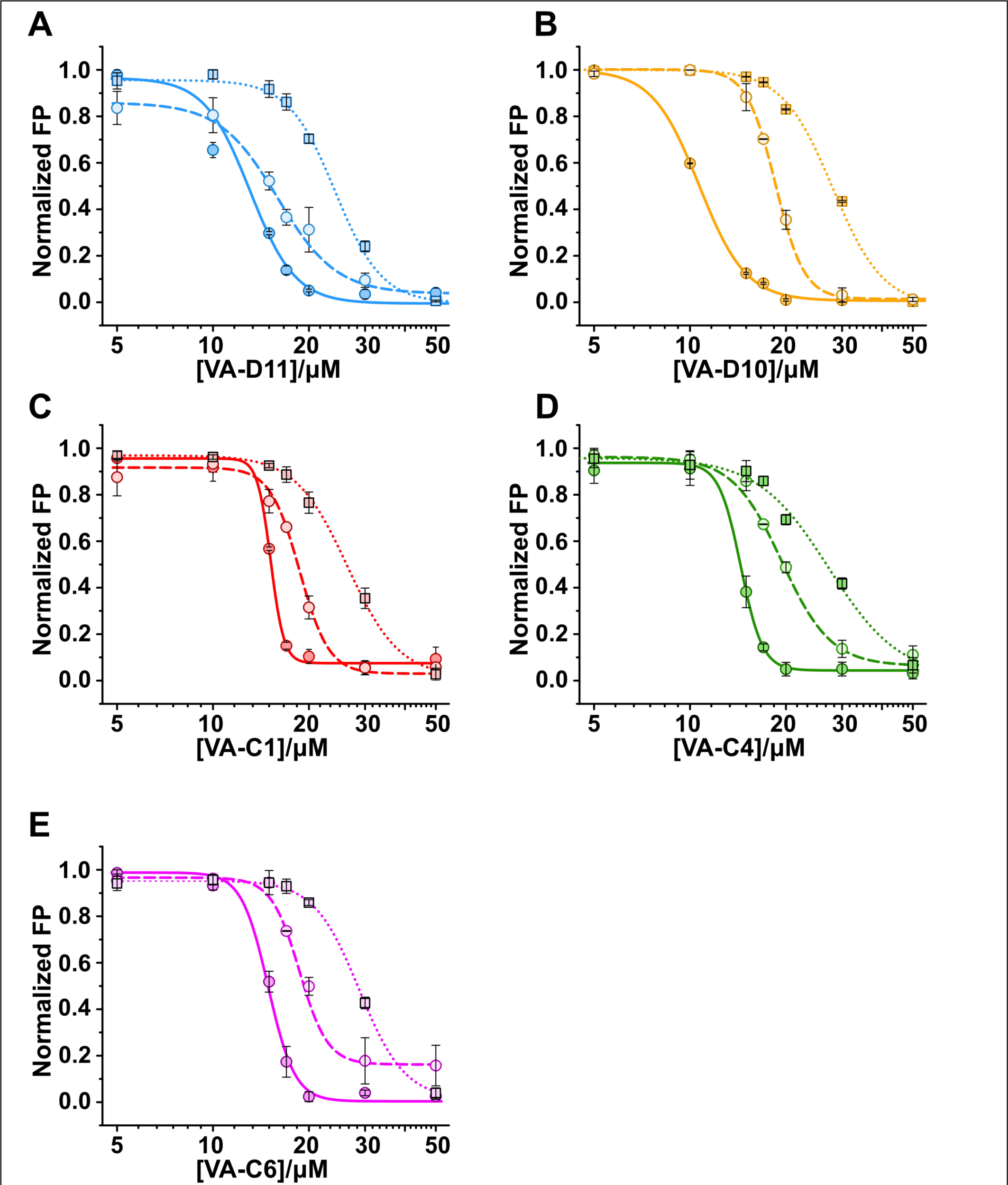
VDAC1 mutants affect VA molecule binding. VA-induced NADH Fluorescence polarization displacement curves from mVDAC1 wild-type (dashed lines) and stabilizing (mVDAC1 N238K - solid line) or destabilizing (mVDAC1 K236A - dotted lines) mutants with increasing concentrations of VA-molecules. K236A and N238K shows decreased and increased affinity, respectively compared to the wild-type channel (KDVA-D11KA=24.2±0.56, KDVA-D10KA =28.0±0.74, KDVA-C1KA =26.4±0.41, KDVA-C4KA =27.22±1.51, KDVA-C6KA =28.8±0.2; KDVA-D11NK=13.1±0.4, KDVA-D10NK =10.7±0.06, KDVA-C1NK=15.2±0.04, KDVA-C4NK =14.4±0.21, KDVA-C6 NK =14.9±0.48). Experiments were performed using a final protein concentration of 7 µM. Data are expressed as mean ± SEM (n = 3).

K236A and N238K, indeed, show decreased and increased affinity, respectively, for VDAC1 compared to the wild-type channel (Table 2). Notably, the NADH affinity for the mutants did not change (Fig. S2), suggesting that residues K236 and N238, previously identified as part of the NADH binding site(Bohm et al., 2020), are only mildly involved in NADH binding. This can be explained by the high mobility of the NADH molecule resolved inside the pore in the NMR (Bohm et al., 2020) structure where the only well-defined region of the NADH binding pocket consists in the nicotinamide moiety anchoring point, but certainly involves several other residues.

**Table 2.** Kd values in NADH - VA-molecules competition assay. In table are reported the Kd values (expressed in µM) obtain after FP assays. Stabilized and destabilized mutant (N238K and K236A) show a variation in affinity with VA-molecules compere to wilt-type protein, increase or decrease respectively. Data are espresse ad mean ± SEM of n=3 independent experiments.

| *Kd [μM]* | *WT* | *N238K* | *K236A* |
| --- | --- | --- | --- |
| **VA-D11** | 15.9 ± 0.7 | 13.1 ± 0.4 | 24.2 ± 0.6 |
| **VA-D10** | 18.5 ± 0.2 | 10.7 ± 0.1 | 28.0 ± 0.7 |
| **VA-C1** | 18.7 ± 0.3 | 15.2 ± 0.1 | 26.4 ± 0.4 |
| **VA-C4** | 19.4 ± 0.4 | 14.4 ± 0.2 | 27.2 ± 1.5 |
| **VA-C6** | 18.7 ± 0.6 | 14.9 ± 0.5 | 28.8 ± 0.2 |

VA molecules, instead, have less steric hindrance and are heavily affected by the mutations, suggesting that the predicted binding pocket is localized within the framing residues that we characterized.

Our data demonstrate that VA molecules bind in a small, well-defined, VDAC1 pocket comprising β-strands 14-16 and the distal part of the N-terminal helix and does not involve the highly mobile residue K20, which is important for NADH binding(Bohm et al., 2020). The SC18 molecule, a recent discovery referenced in literature(Heslop et al., 2022), shares a similar region with VA molecules, although its precise binding site remains undefined. In contrast to SC18, VA molecules are marginally larger but lack nitro groups, which are generally considered toxic (Nepali et al., 2019).

### VA molecules selectively kill cancer cells

The small molecular weight of the VA molecules, the satisfaction of the Lipinski’s rule of five (Lipinski et al., 2001), together with their ability to interfere with VDAC1 NADH binding and N-terminal helix movement, prompted us to test whether VA molecules can be effectively used to halt cancer cell proliferation.

We initially tested the five selected molecules on the highly aggressive, highly glycolytic, breast cancer cell line SKBR3. All VA molecules show a marked reduction (∼75% at VA high dose) in cell viability after 48h of treatment. In the same set of experiments, the non-tumorigenic fibroblast cell line NIH-3T3 was not affected by the treatment (Fig. 6A-E). Notably, SKBR3 cells show increased level of VDAC1 protein expression compared to NIH-3T3 (Fig. S3A). The calculated EC_50_ aligns with the K_D_ values found on the binding assays performed on the purified VDAC1 (Fig. 6F), confirming that the effect is specific to VDAC1.

**Figure 6.**
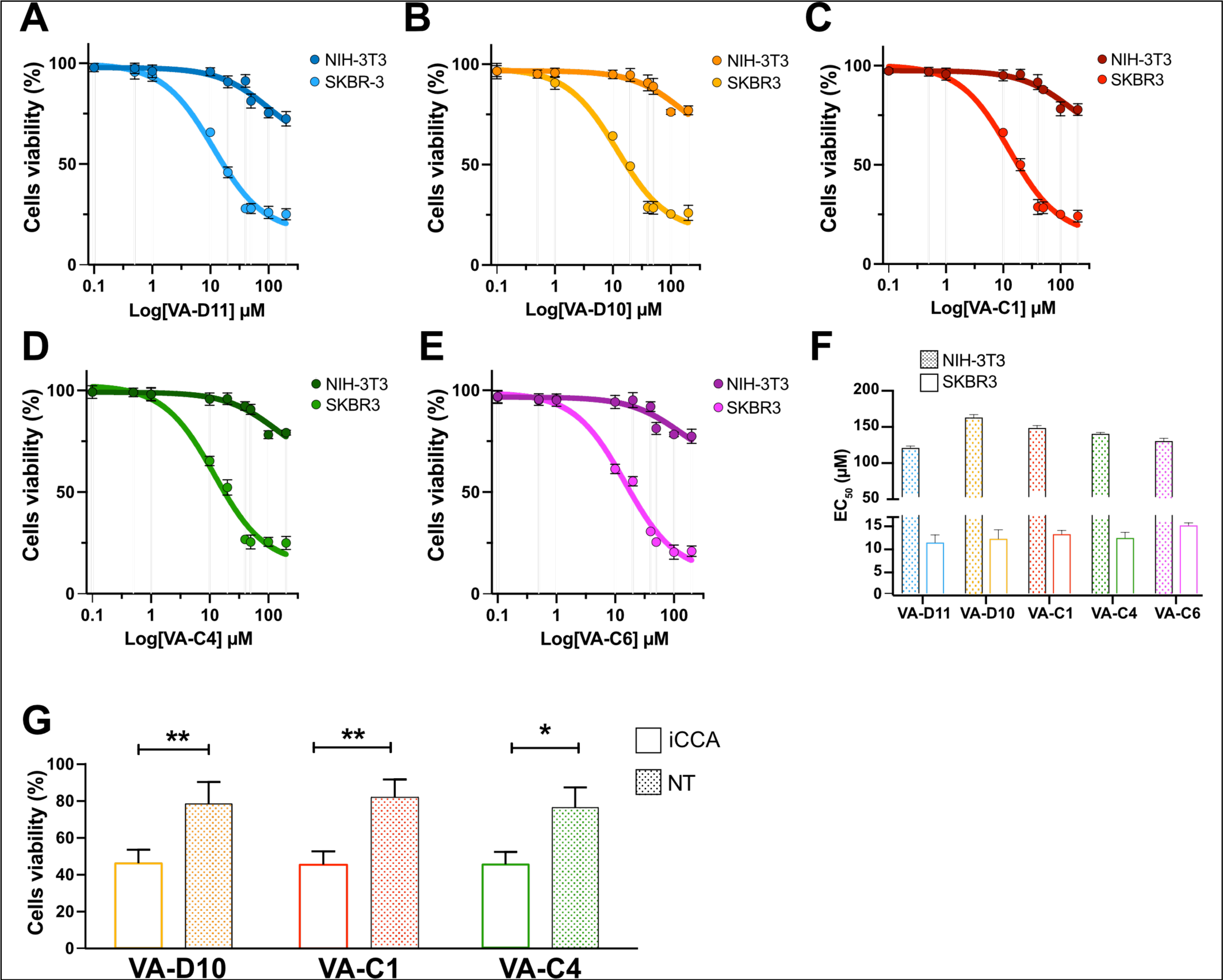
VA molecules affect the viability of cancer cells. (A-B-C-F-E) Dose-response curves representing cell viability of NIH-3T3 and SKBR-3 treated with different concentrations of VA molecules (VA-D11, VA-D10, VA-C1, VA-C4, and VA-C6). Data are expressed as mean ± SEM (n = 3). (F) Bar plot reporting Half the effective dose (EC50) retrieved after cell viability assay. SKBR-3 cell lines show greater sensitivity to treatment with the five molecules (VA-D11: 10.01 ± 1.51 µM; VA-D10 10.22 ± 1.45 µM; VA-C1 10.93 ± 1.88 µM; VA-C4 10.83 ± 2.87 µM; VA-C6 10.38 ±2.19 µM) then NIH-3T3 cell lines (VA-D11: 118.1 ± 2.79 µM; VA-D10 160.3 ± 3.98 µM; VA-C1 145.8 ± 3.22 µM; VA-C4 137.5 ± 2.11µM; VA-C6 127.5 ± 3.92 µM). (G) The effect of VA molecules (D10, C1, C4) on the viability of iCCA patient-derived and paired non-tumor (NT) 2D cells. Viability was measured by MTT assay after 72h exposure using 2μM concentration (*p<0,05,**p<0,01). Data are represented as mean ± SEM (n=3).

SKBR3 and NIH-3T3 are, however, commercial immortalized cell lines that hardly reproduce the tumor environment. To validate the effectiveness of the VA molecules in potentially treating cancer, we tested the molecules on primary cells derived from intrahepatic cholangiocarcinoma patients (iCCA) and on paired non-tumor liver cells, having a different expression level of VDAC1 (Fig. S3B). Out of the five molecules, VA-C1, VA-D10 and VA-C4 show a similar reduction in iCCA cell viability of about 50% after 72h of exposure to 2μM concentration of molecules (Fig. 6G). Again, control cells were unaffected by the treatment (Fig. 6G), underlining the ability of VA molecules to kill the cancer cells.

### VDAC1-mediated disruption of NADH balance impairs mitochondrial function in glycolytic tumors

Given that (1) NADH crosses the outer mitochondrial membrane through VDAC1 (Shoshan-Barmatz et al., 2010), (2) VDAC1 is overexpressed in many tumor cells lines(Wang et al., 2022) including iCCA and SKBR3 (Fig. S3) and (3) the SC18 molecule targeting the VDAC1 NADH binding site(Heslop et al., 2022) reduces mitochondrial NADH, we hypothesized that VA molecules can similarly impact mitochondrial energy generation by blocking the NADH transport. To test the hypothesis, we analyzed the oxygen consumption rate linked to oxidative phosphorylation (OXPHOS) in SKBR3 and NIH-3T3 cells by high-resolution respirometry (HRR). Fig. S6 shows a representative respirometry curve of untreated NIH-3T3 and the specific protocol used. We first analyzed the OXPHOS respiration relative to the NADH electron transport pathway (N-pathway) driven by Complex I. This state was achieved in the presence of a combination of NADH-generating substrates (pyruvate, malate and glutamate) and saturating concentration of ADP, after a mild permeabilization of the plasma membranes. Then, the addition of succinate allowed the activation of Complex II and the measurement of the succinate pathway in combination with the N-pathway (NS-pathway).

Oxygen consumption, corresponding to the reported respiratory states in cells treated with 10 µM of VA molecules for 48 h, was monitored and compared to the control. As shown (Fig. 7A and Table S2), a dramatic reduction of oxygen flows was detected in both N-Pathway and NS-Pathway, ranging from 15% to 40% in tumoral cell lines. Otherwise, in the same set of experiments, the NIH-3T3 cells were unaffected by the small molecule treatment. According to previous data, the net OXPHOS fluxes, corresponding to the oxygen consumption directly devoted to the ADP phosphorylation, were similarly reduced (Fig. 7B).

**Figure 7.**
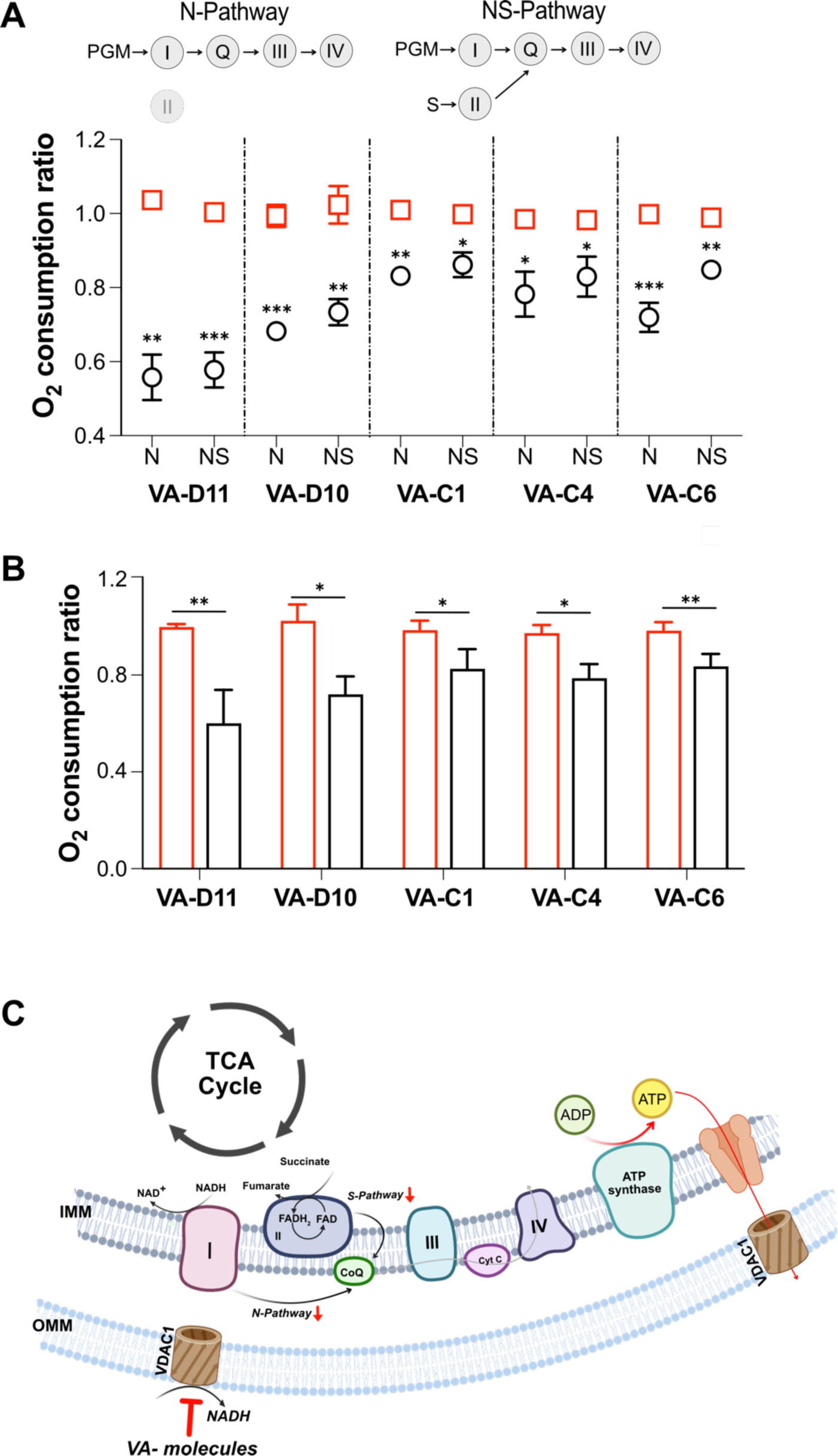
High-resolution respirometry shows cancer cell mitochondrial distress upon VA treatment. (A) Quantitative analysis of the oxygen consumption in the analyzed states (N-Pathway and NS-Pathway) of SKBR-3 and NIH-3T3 attained after the exposure to 10 µM VA-molecules for 48h. Data are indicated as the mean ± SEM (n = 3) and expressed as the ratio of the treated cells to the untreated ones. Data were analyzed by unpaired t-test, with *p < 0.05, **p < 0.01 and ***p < 0.001. (B) Quantitative analysis of the net oxygen consumption in the NS-Pathway or ATP-linked flux of SKBR-3 and NIH-3T3 attained after the exposure to 10 µM VA-molecules for 48h. Data are indicated as the mean ± SEM (n = 3) and expressed as the ratio of the treated cells to the untreated ones. Data were analyzed by unpaired t-test, with *p < 0.05, **p < 0.01 and ***p < 0.001. (C) Proposed mechanism for NADH imbalance through VDAC1.

Overall, these results indicate that the VA molecules can affect mitochondrial functionality only in high glycolytic cell lines, confirming there use as a potential therapeutic intervention for cancer treatment.

### Establishment of organoid cultures for VA molecules testing

In order to verify the efficiency of VA molecules treatment in a reliable tumor-like context, we developed a suitable three-dimensional system of organoids from patients undergoing diagnostic surgical resection for iCCA.

Organoids were obtained from both iCCA and non-tumor tissues in 5 of the 6 patients included in the study, considering our cohort’s underlying disease spectrum (Table S3).

Tumor organoids allowed long-term *in vitro* expansion, in contrast with organoids derived from non-tumor tissues, which showed a more robust proliferation at the early stage of culture, but after a few months stopped to proliferate. Organoids morphologically showed a cystic structure, with an external layer that defined an internal core (Fig. S4A), features that have described (Saito et al., 2019).

Further, we characterized the organoids by immunofluorescence technique to map the presence of molecular markers, attesting their reliability for drug testing. We revealed the expression of typical biliary markers, such as Epithelial Cell Adhesion Molecule (EpCaM), cytokeratin 7 (CK7), and cytokeratin 19 (CK19) (Fig. S4B). Moreover, we analyzed the level of nuclear antigen Ki67 in tumor organoids, confirming that its presence >5% is an important requirement for successful organoid derivation (Fig. S4B) (Nuciforo et al., 2018).

RT-PCR performed on iCCA cells compared to non-tumor cells showed overexpression of VDAC1 mRNA exclusively in cancer cells (Fig. S3B), leading us to use this system to test the effect of VA molecules on 3D cultures.

### Organoids are valid pre-clinical models to test VA molecules efficacy

The absence of appropriate *in vitro* models that accurately represent the cancer biology and heterogeneity of patients is a significant barrier to the development of new therapies and biomarkers for iCCA. Due to their ability to retain the morphological properties of primary tissue and assess different biological features, including proliferation, signaling, and cell-cell interaction, liver organoids could represent an ideal approach (De Siervi and Turato, 2023).

Based on this premise, we tested the impact of different dosages of VA molecules both in iCCA and non-tumor organoid cultures, monitoring cell viability after 72h. In these *in vitro* assays, we observed that VA molecules treatment selectively decreased iCCA-derived organoids growth in a dose-dependent manner, with an EC_50_ of 24.7±5.9, 14.6±1.8 and 18.7±1.1 μM for VA-D10, VA-C1 and VA-C4, respectively (Fig. 8). Strikingly we measured a very low effect on the viability of non-tumor cells. In addition, we compared *in vitro* VA molecules activity with the gemcitabine (GMB), clinically used for iCCA treatment. GMB demonstrates significant efficacy against both iCCA and control organoids, even at concentrations five times lower than those of VA molecules (Fig. S5). This underscores the enhanced chemical potency of VA molecules compared to standard drugs.

**Figure 8.**
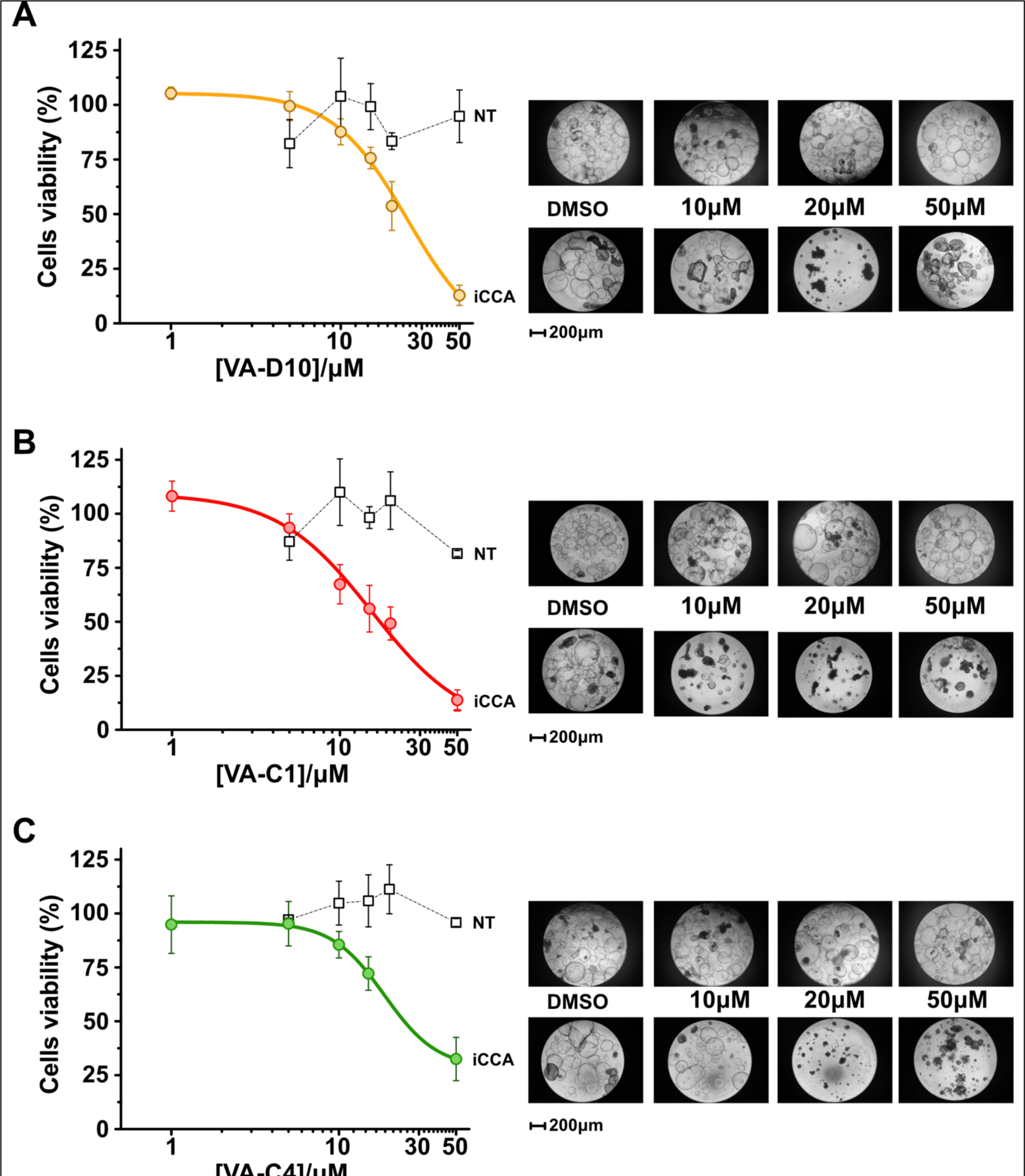
Cholangiocarcinoma 3D models are affected by VA treatment. Tumor (iCCA) and non-tumor (NT) organoids were exposed to VA treatment for 72h at the indicated concentration. DMSO-treated organoids were used as control. (A-C) Left panel: VA treatment (D10, C1, C4) reduce cell viability of iCCA organoids in a dose-dependent manner, while show a limited effect on non-tumor organoids. Data are represented as the percentage of control DMSO-treated organoids and are the mean of at least three independent experiments ± SEM. Right panel: representative bright-field images of tumor and non-tumor organoids exposed to VA treatment (D10, C1, C4) at the indicated concentration for 72h. Scale bar: 200 μm.

Taken together, these results demonstrate that selectively displacing NADH from VDAC1 cavity hampers cancer cell survival and show the first evidence that novel chemical entities specifically targeting the major mitochondrial porin are effective on patients’ derived tumor primary cells, paving the way for the development of precision medicine VDAC1-targeted drugs.

## Discussion

Small molecule modulation of cell fate is one of the major research areas in cancer biology, but it is still one of the most challenging. Cancer research, indeed, must face the heterogeneity of cancer phenotypes, the individual immune response to the disease (De Palma and Hanahan, 2012) and the general cytotoxicity of chemotherapeutics (Fu et al., 2018). These translate in efforts to develop specific anticancer agents. The long process from bench research to clinical trials additionally slows down the generation of novel classes of molecules.

Targeting cancer metabolism as a universal strategy has been proposed for years. However, most of the efforts are directed towards either the inhibition of enzymes involved in the glycolysis (Pelicano et al., 2006) or impairing the mitochondrial activity (Zong et al., 2016), relying on the high glycolytic activity of certain cancer types or in the mitochondrial dysfunction on others. A unique molecule capable of playing on both sides is still elusive.

Rather than just a mere channel, VDAC1 figures as a molecular switch for the modulation of the cell bioenergetics and links glycolysis to mitochondrial metabolism by its direct interaction with HKs. Except for some compensatory effect given by other isoforms, the inactivation of VDAC1 gene triggers a re-arrangement of the electron transport system functioning (Magri et al., 2023a). Where VDAC1 represents the only functional porin, as in yeast, its deletion even prompts the mitochondrial switch off and a complete metabolic rewiring aimed at bypassing the organelle dysfunction (Magri et al., 2020). Therefore, targeting the major mitochondrial metabolite transporter VDAC1 is a promising strategy since the channel is involved in many cancer processes, i.e. the inhibition of apoptosis, and it is considered the ‘mitochondrial gate keeper’ because of its role in metabolite translocation from the cytosol to the mitochondria and vice-versa. However, most of the identified molecules are either not specific, influence the channel activity, or their binding pocket has not been fully characterized (Ben-Hail et al., 2016; Magri et al., 2018), making any effort harder for a further pharmaceutical development. A number of agents have been employed to disrupt the HK-VDAC1 complex (clotrimazole or 3-bromopyruvate) (Krasnov et al., 2013), inhibit VDAC1 conductance (DIDS and derivatives) (Ben-Hail and Shoshan-Barmatz, 2016) or reduce VDAC1 expression (Reina and De Pinto, 2017). However, many of them could affect cancer cells and normal tissues, especially those that use glucose as the main energy source (Shoshan-Barmatz et al., 2017).

In light of these considerations, we rationalized that a pocket further away from VDAC1 central cavity would be the ideal candidate to search for. This region seems to be shared by the recently discovered SC18 molecule (Heslop et al., 2022) with some differences, even though its exact binding pocket has not been uniquely identified. Following this line, we developed a workflow for identifying and testing small molecules targeting VDAC1 using a combination of direct binding detection on the purified target and further validation on patient-derived primary cell cancer organoids. Using a bottle-neck strategy, we identified three drug-like candidates that perform well on cholangiocarcinoma organoids and adherent cancer cell models, but at the same time they leave unperturbed healthy cells and organoids. VA molecules are slightly bigger than SC18, but on the contrary they do not present nitro-groups, which are often considered toxic (Nepali et al., 2019). Compared to other synthetic molecules targeting VDAC1 (i.e. VBIT-3/4) (Ben-Hail et al., 2016), VA molecules are significantly smaller (average number of atoms is 39 and 52 for VA and VBIT molecules, respectively). We further uniquely identified their binding site within VDAC1 and discovered that our small molecules, at micromolar concentrations, lead to the displacement of NADH from its anchor point to the protein. The NAD^+^/NADH homeostasis is a key player in cancer cell survival because of its role in metabolic advantage for sustained cell proliferation (Moreira et al., 2016). The NAD^+^ depletion through mutations in the isocitrate dehydrogenase genes has been shown to dramatically affect the mitochondrial TCA cycle and solid tumor vitality (Tateishi et al., 2015). TCA cycle metabolites themselves are physiological signaling molecules (Martinez-Reyes and Chandel, 2020) able to control anabolism, catabolism and cell fate. In highly glycolytic cancer cells (such as iCCA or the commercial breast cancer SKBR3 used in this study), the altered mitochondrial bioenergetics lead to uncontrolled proliferation and tumor progression. High lactate production, indeed, is oxidized back to pyruvate to foster TCA cycle. Therefore, a higher demand for NADH is required for the TCA cycle to proceed. Hence, limiting the NADH availability can slow down the mitochondrial electron transport capacity by directly influencing the Complex I (NADH ubiquinone-oxidoreductase) and indirectly the Complex II (Succinate dehydrogenase) by limiting its own substrate.

As a matter of fact, HRR data showed a tremendous impact on mitochondrial respiration in our cancer cell model following treatment with molecules and, precisely, in the measurement achieved in the exclusive presence of NADH-linked substrates. This data suggests a significant reduction of the Complex I contribution to the OXPHOS respiration, possibly due to a limited amount of NADH, which reaches mitochondria through VDAC1. The impact of VA molecules on respiration appears even more drastic considering that the further activation of Complex II by succinate cannot overcome the mitochondrial dysfunction. Indeed, the oxygen flows relative to the total OXPHOS respiration (NS-pathway and the ATP-related fluxes) are significantly lower in VA treated cells compared to the untreated controls. Therefore, it could be hypothesized that the binding of these drugs would lead to a change in the translocation of NADH or other mitochondrial metabolites through the porin, causing an imbalance in the TCA cycle by limiting the cofactors necessary for the first respiratory chain reaction complex in which mitochondria respiration are affected.

The results are strengthened by the null effect of VA molecules on non-tumor cell lines. The expression of VDAC1 is increased in numerous human cancer cell lines compared to normal cell lines (Shinohara et al., 2000; Shoshan-Barmatz and Ben-Hail, 2012; Simamura et al., 2006; Simamura et al., 2008), and this could explain the different impact on mitochondrial activity and cell proliferation observed. The strategy of using patient-derived organoids has been successful in recognizing only the small molecules that potentially would be effective *in vivo*, by excluding (1) some of the cell biology hits that were not effective on fibroblasts but were killing the organoids and (2) all the compounds that were also killing the control organoids.

The identified compounds have an average EC_50_ in the micromolar range, therefore they cannot be directly used as therapeutics. However, we (1) have identified a precise hot-spot in VDAC1 that could be an easy target for additional drug-screening campaigns; (2) isolated three effective drug-like novel chemical entities that can be further optimized by simple medicinal chemistry and biochemical studies as reported; (3) designed a simple killing method using organoids that will boost the further drug development of these compounds since the animal models are not any longer required to proceed to clinical trials if scaffolded 3D cultures are used (Wadman, 2023).

## Methods

### In silico screening

Commercially available libraries of millions of chemical compounds (MCule (10M) [*mcule database*. https://mcule.com/database/] and Enamine in-stock (2.5M) [https://enamine.net/compound-collections/screening-collection] have been screened by Atomwise using its deep learning-based computational drug discovery platform (AtomNet^TM^)(Stahl et al., 2018; Wallach et al., 2015; Wallach and Heifets, 2018). After rescoring and selection, 70 compounds were purchased from Enamine and tested experimentally.

Docking simulations have been performed using the free software *AutodockVina* (Eberhardt et al., 2021) with the mouse VDAC1 crystal structure (PDB: 3EMN) as target protein and the 3D structures of VA molecules generated with *Open Babel* (O’Boyle et al., 2011) as flexible ligands.

### Protein expression and purification

Mouse VDAC1 (mVDAC1) wild-type and mutant constructs were expressed as reported in (Ujwal et al., 2009). Briefly, mVDAC1 and mutants cloned in pQE9 vector were transformed into M15 *E. coli* for protein expression. Cells were grown at 37°C in LB medium until OD_600_= 0.6 and induced for 4 h with 1 mM IPTG. Cells were harvested and resuspended in lysis buffer (50 mM Tris-HCl (pH 8.0), 2 mM EDTA, and 20% sucrose). The resuspended pellet was sonicated and centrifuged (12.000 x g, 20 min) to obtain inclusion bodies. The inclusion body pellet was washed with wash buffer (300 mM NaCl, 20 mM Tris-HCl (pH 8.0), 2 mM CaCl_2_) and solubilized in equilibration buffer (300 mM NaCl, 20 mM Tris-HCl (pH 8.0), 6 M guanidinium hydrochloride) for 3h at 4°C under stirring. Solubilized inclusion bodies were applied to the HisTrap HF 5 mL affinity column (Cytiva), washed with equilibration buffer containing 30 mM imidazole (5CV), and eluted with equilibration buffer containing 150 mM imidazole (8CV).

*In vitro* refolding was performed by dropwise dilution (100 µL/min) of solubilized protein into refolding buffer (300 mM NaCl, 1 mM EDTA, 5 mM DDT, 25 mM Tris-HCl (pH 8.0) and 1% LDAO) (mg protein:ml buffer = 1:20). The procedure was carried out at 4°C for 3h. Subsequently, the refolded protein was dialyzed against dialysis buffer (300 mM NaCl and Tris-HCl pH 8.0) at 4°C overnight. To reduce detergent concentration, the refolded protein was applied to HisTrap HP 5ml affinity column, washed with equilibration buffer (300 mM NaCl, 25 mM Tris-HCl (pH 8.0), 0.1% LDAO) containing 30 mM imidazole (5CV), and eluted with equilibration buffer containing 250 mM imidazole (8CV). Refolded protein was concentrated by using Amicon Ultra-30KDa and applied to a Superdex 200 increase 10/300 column (Cytiva) and eluted with SEC buffer (50 mM NaCl, 20 mM Tris-HCl (pH 8.0), 0.1% LDAO). Protein purity and identity were verified by SDS/PAGE.

### NMR spectroscopy

For validation by NMR spectroscopy, each compound was dissolved in d6-DMSO at a final concentration of 500 µM. ^1^H NMR spectra were recorded at 310 K at a Bruker 700 MHz spectrometer equipped with a TXI probe, with an acquisition time of 2.36 s, inter-scan delay of 8 s, 4 dummy scans and 32 scans. The spectra were processed in Topspin by applying gaussian broadening (GB 0.02, LB-0.1 Hz) and baseline corrected. ^19^F NMR spectra were recorded on the same samples at a Bruker 800 MHz spectrometer equipped with a TXI probe with the ^1^H channel tuned to the ^19^F Larmor frequency at 752.9 MHz. The spectra were recorded at 298 K with an acquisition time of 0.56 s, inter-scan delay of 5 s, 0 dummy scans and 256 scans, and processed by applying exponential broadening (LB 2 Hz). For protein-ligand interaction experiments, each compound was dissolved at a final concentration of 100 µM in 20 mM Tris HCl buffer, pH 8.0, 50 mM NaCl, 0.1% LDAO either in the absence or in the presence of 5 µM VDAC1 diluted from a 115 µM stock solution in the same buffer. ^19^F NMR spectra were recorded at 298 K with an acquisition time of 0.56 s, inter-scan delay of 1 s, 0 dummy scans and 256 scans, and processed by applying exponential broadening (LB 2 Hz).

### Fluorescence Polarization Assay

All Fluorescence Polarization measurements were performed on a FluoStar-Omega microplate reader (BMG LABTECH) in black opaque 96-well microplates (Costar #3925) with excitation and emission wavelengths of 482 nm and 530 nm, respectively. Fluorescence polarization assay was carried out by titrating mVDAC1 wild-type and mutants (N238K and K236A) with 5 µM NADH. The fluorescence polarization using the following equation:

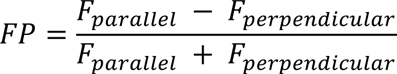

where FP is the fluorescence polarization reading, F-parallel is the fluorescence intensity parallel to the excitation plane, and F-perpendicular is the fluorescence intensity perpendicular to the excitation plane. The dissociation constant (K_D_) was obtained by fitting the fluorescence polarization values and the corresponding protein concentrations into a nonlinear regression model in Origin Lab. For the FP competition assays, different concentrations of VA molecules were titrated against 7 µM of VDAC1 and 5 µM of NADH. The K_D_ was obtained by fitting the fluorescence polarization values and the corresponding VA molecules concentrations into a Hille curve. All experiments were performed in triplicates using 100 μL per well in 50 mM NaCl, 20 mM Tris (pH 8.0), and 0.1% w/v LDAO.

### Thermostability Assay

For the thermal denaturation assay, mVDAC1 and mutants (at a final concentration of 0.9 mg/ml) in SEC buffer were pre-treated with 1 mM of VA molecules or 0.5 % DMF (as control) for 1h at 4°C. Subsequently, samples were loaded into quartz capillaries (Nano-Temper Technologies, Germany), and experiments were carried out using the NanoTemper Tycho NT.6 (NanoTemper Technologies, Germany). The temperature gradient was set to an increase of 30 °C/min from 35° C to 95° C. Protein unfolding was measured by detecting the temperature-dependent change in tryptophan fluorescence at emission wavelengths of 330 and 350 nm. Inflection temperatures (T_i_) were determined by detecting the maximum of the first derivative of the fluorescence ratios (F350/F330), using the internally automated evaluation features of the NT.6 instrument. The variation of T_i_between treated (T_iT_) and untreated (T_iU_) samples was calculated by the formula ΔT_i_= T_iU_-T_iT_. To quantify the effect of mVDAC1 mutations in modifying VA molecules’ stability, ΔΔT_i_ was calculated with the formula ΔΔT_i_= ΔT_iWT_ - ΔT_iMut_.

Positive values of ΔΔT_i_show the stabilizing influence of mutations, while negative values denote the destabilizing effect of mutations. All measurements were repeated at least three times.

### Cell cultures and viability assays

SKBR-3 (Cat # HTB-30) and NIH-3T3 (Cat # CRL-1658), were purchased from the American Tissue Culture Collection (ATCC). SKBR-3 cells were grown in McCoy’s 5A Medium supplemented with 10% fetal bovine serum (FBS), 100 units/mL penicillin and 100 μg/mL streptomycin. NIH-3T3 cells were grown in Dulbecco’s Modified Eagle’s Medium (DMEM) high glucose with the addition of 10% Calf Bovine Serum (CBS), 100 units/mL penicillin and 100 μg/ mL streptomycin. Both cell lines were maintained in 5% CO_2_/air at 37 °C. All experiments were performed when cells reached 70– 80% confluency. Each cell line was treated with different VA molecules concentration for 48 h. Subsequently, the cells were used for cell viability assay.

Cell viability was evaluated by 3-(4,5-dimethyl thiazolyl-2)-2,5-diphenyl tetrazolium bromide (MTT) assay. SKBR-3 and NIH-3T3 cell lines were plated in 96-well plates at 10.000 cell/well and after 24 h received the indicated treatment. Subsequently, MMT was added to the culture medium to reach a final concentration of 5 mg/ml. After incubation at 37°C for 3 h, the medium was removed, and the formazan crystals produced were dissolved by adding 100 µl of dimethyl sulfoxide. The absorbance at 590 nm was determined using the microplates reader Varioskan (Thermo Scientific).

### High-resolution respirometry

The characterization of mitochondrial respiration in the SKBR-3 and NIH-3T3 cell lines was performed by High-Resolution Respirometry using the two-chamber system O2k-FluoRespirometer (Oroboros Instruments, Innsbruck, Austria). The whole respiratory profiles of SKBR-3 and NIH 3T3 cell lines, exposed or not to 10μM of small molecules for 48 h, were determined by a specific substrate-uncoupler-inhibitor titration (SUIT) protocol as in (Leggio et al., 2022) in mitochondrial respiration buffer Mir05 (Oroboros Instruments) at 37 °C under constant stirring of 750 rpm.

Plasma membranes of intact cells were mildly permeabilized with 3μM digitonin and the non-phosphorylating respiration (LEAK) was measured in the absence of adenylates. Oxygen flow-related N-pathway was attained by the addition of pyruvate (5 mM), glutamate (10 mM), malate (2 mM), and ADP (2.5 mM). NS-pathway was then activated by supplementation with 10 mM succinate. The maximal capacity of electron transport (ET) chain was determined by titration with the uncoupler carbonyl cyanide 3-chlorophenylhydrazone (CCCP, 0.5 μM) up to the complete dissipation of the proton gradient. Finally, the residual oxygen consumption (ROX) was achieved by inhibiting the ET chain with 2 μM rotenone and 2.5 μM antimycin.

Oxygen fluxes corresponding to LEAK, N-, NS-pathway and ET maximal capacity were normalized for the ROX and expressed as pmol/s per million cells (Table S3). The oxygen flows linked to ATP production were calculated as the difference between the NS-pathway and the LEAK respiration(Magri et al., 2023b). Respiratory states of cells exposed to VA molecules were normalized to their respective untreated controls and shown as mean ± SEM of n=3 three independent experiments. Data were statistically analyzed by unpaired t-test using Prism software (GraphPad, San Diego, CA, co USA). The following values * p < 0.05, ** p < 0.01, *** p < 0.001 were taken as significant.

Instrumental and chemical background fluxes were calibrated as a function of the oxygen concentration using DatLab software (Oroboros Instruments). All substrates were purchased by Sigma Aldrich.

### Western blots

1 x 10^6^ cells for each line were collected and lysed in extraction buffer (50 mM Tris pH=7.4, 150 mM NaCl, EDTA 1 mM, 1% TRITON X-100 and protease inhibitors). After lysis, the supernatant was collected, and the protein content was estimated with the Bradford reagent with bovine serum albumin (BSA) as a standard. 50 µg of total protein of each sample were separated using SDS/PAGE electrophoresis, transferred to a nitrocellulose membrane (AmershamProtran 0.45 µM NC; GE Healthcare Life Sciences), and blocked in 5 % BSA at room temperature for 1h. The membranes were incubated overnight at 4 °C with primary antibodies against VDAC1 and β-Actin (1:5000, #2146; Cell Signaling Technology) after the washing step with secondary antibodies HRP-conjugate Goat anti-Mouse IgG (H+L) (1:5000, #89842, Invitrogen), HRP-conjugate Goat anti-Rabbit IgG (H+L) (1:5000, #31460, Invitrogen). Chemiluminescent detection was performed using Pierce™ ECL Western Blotting Substrate. The membranes were read using the Azure c600 Imaging System (Azure Biosystems). Data were analyzed with Image-StudioLite software (Li-Cor Biosciences), and β-Actin was used as an internal loading control. Data were statistically analyzed by one-way ANOVA followed by Dunnett’s multiple comparisons test.

### Primary cell culture and organoid generation

For organoid generation, surgically resected iCCA specimens along with matched non-tumoral tissue from a non-adjacent liver site were obtained from treatment-naïve patients at Fondazione IRCCS Policlinico San Matteo, Pavia, Italy. Tissue samples were stored in tissue storage solution (Miltenyi Biotec, Bergisch Gladbach, Germany) and processed within 24 hours. A written informed consent was obtained from each individual. The study protocol is compliant with the ethical guidelines of the 1975 Declaration of Helsinki and was approved by our institutional ethical committee. Briefly, tissue samples were treated by enzymatic and mechanical dissociation with the human Tumor Dissociation Kit and gentle MACS Dissociator, according to the manufacturer’s instructions (Miltenyi Biotec). The cell suspension was filtered in a 70 µm cell strainer and centrifuged twice at 50 g for 2 min. The cell pellet was then incubated in 1ml of ACK lysing buffer (GIBCO) to remove blood cells. After cell counting, about 15.000 cells per well were resuspended in Cultrex UltiMatrix Reduced Growth Factor (RGF) Basement Membrane Extract (BME) (R&D Systems) and seeded in pre-heated low adhesion 24-well plates (GreinerBio). The remaining cells were cultured in Dulbecco’s Modified Eagle’s Medium (DMEM)/F-12 (Sigma-Aldrich) with 10% Fetal Bovine Serum (FBS) (GIBCO), and 1% penicillin/streptomycin (GIBCO) and seeded in T25 flask, in order to generate bidimensional primary cell lines. After BME droplets polymerization at 37°C, an expansion medium was added. The expansion medium was adapted from the protocol of Nuciforo et al (2018)(Nuciforo et al., 2018) and composed of Advanced DMEM/F-12 (GIBCO) with 1% penicillin/streptomycin (GIBCO), 1% Glutamax (GIBCO), 10 mM HEPES (GIBCO) with the addition of B-27 (1:50) (GIBCO), N-2 (1:100) (GIBCO), 10 mM nicotinamide (Sigma-Aldrich), 1.25 mM N-acetyl-L-cysteine (Sigma-Aldrich), 10 µM forskolin (Tocris), 10 µM ROCK inhibitor (Y-27632) (Tocris), 5 µM A83-01 (Sigma-Aldrich), 10 nM [Leu15]-gastrin I (Sigma-Aldrich), 50 ng/ml epidermal growth factor (EGF) (R&D Systems), 25 ng/ml hepatocyte growth factor (HGF) (Homemade), 10% R-Spondin1 conditioned medium (Sigma-Aldrich) and 30% Wnt3a conditioned medium with Noggin (Sigma-Aldrich). The passage of the organoids was performed periodically, every 7-10 days, by incubation in Cultrex Organoid Harvesting Solution (R&D Systems) for 1 hour on ice and mechanical dissociation with a pipette. For freezing, the organoids were dissociated and resuspended in Recovery Cell Culture Freezing Medium (GIBCO) and then cryopreserved.

### Drug Treatment

Tumor and non-tumor organoids were plated in 20 μl of BME droplets in 48-well plates in order to develop organoids. After 3 days, VA small molecules (D10, C1, and C4) at different concentrations were resuspended in expansion medium with the addition of 0.02% of Tween20, and cell viability was measured after 72 hours, using CellTiter-Glo 3D Cell Viability Assay (Promega). Also, tumor and paired-non tumor bidimensional cell lines were seeded at density of 5.000 cells per well in 96-well plates, and after 3 days treated with VA small molecules at different concentrations, and viability was evaluated by MTT assay after 72 hours. Luminescence and absorbance respectively were measured on Polarstar-Omega microplate reader (BMG LABTECH). Results were normalized to vehicle (100% DMSO). Curve fitting was performed using Prism (GraphPad) software and the nonlinear regression equation. All experiments were performed at least three independent times, and results are shown as mean ± SEM.

## Acknowledgments

This work was supported by ‘Programma per Giovani Ricercatori - Rita Levi Montalcini 2016’ granted by the Italian Ministry of Education (Project number: PGR16HTPSF) and by the Department of Molecular Medicine of the University of Pavia, Italy under the initiative Dipartimenti di Eccellenza (2018–2022) to M.L and Immuno-HUB_T4-CN-02-anno 2021 to C.T.

