## Supplemental Figures and Methods for "VDAC1 selective molecules promote patients’-derived cancer organoids death through mitochondrial-dependent metabolic interference"

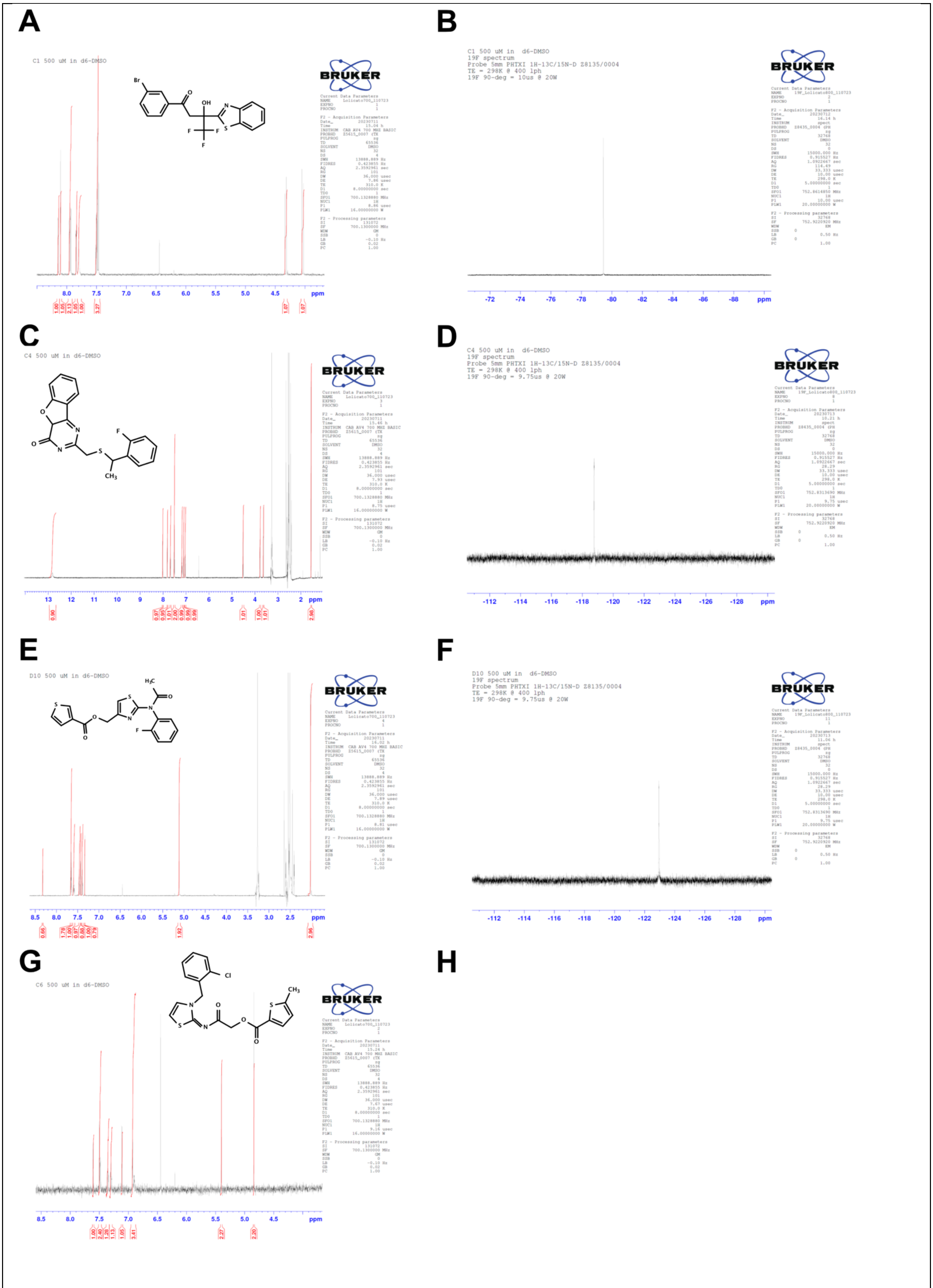


**Figure S1. $^{1}$H and $^{19}$F NMR spectra of VA compounds**
A, C, E and G are the 1H NMR spectra of VA-C1, VA-C4, VA-D10 and VA-C6, respectively. B, D and F are the $^{19}$F spectra of the same compounds.

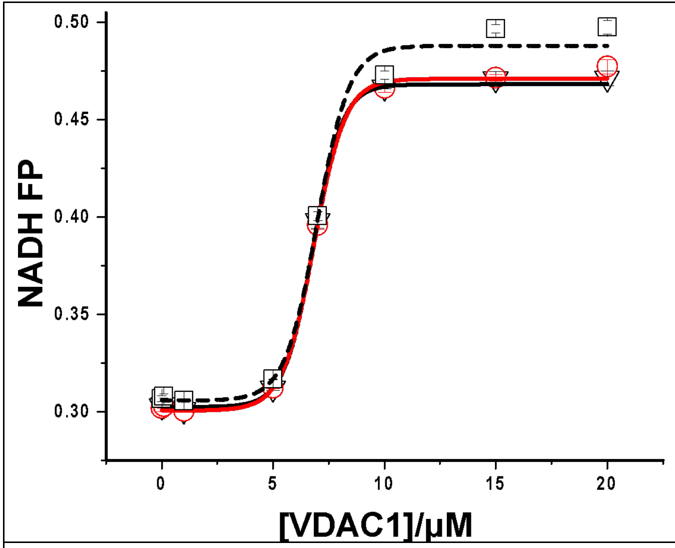


**Figure S2. NADH binding to purified VDAC1 measured by Fluorescence Polarization**
VDAC1 wild-type (triangles) and mutants (K236A, square and N238K, circles) purified protein has been titrated against 5μM NADH and its fluorescence polarization change has been monitored using λex= 482nm and λem= 530nm. Calculated $K_D$ values are (μM): 6.80±0.06, 6.97±0.16 and 6.83±0.06 for VDAC1 wild-type, K236A and N238K respectively.

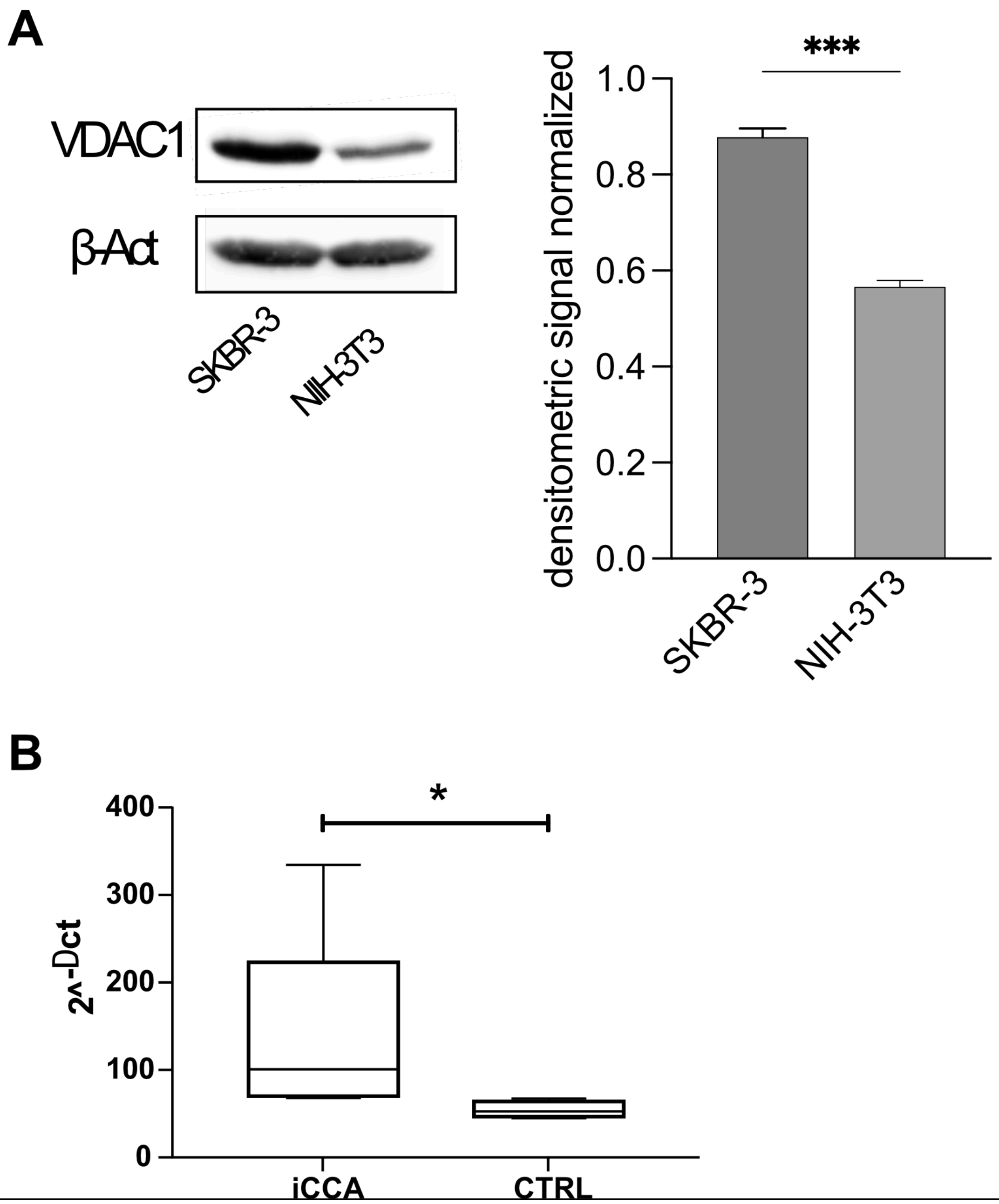


**Figure S3. VDAC1 expression levels on SKBR3 and iCCA**

(A). Western blot illustration and relative densitometric quantification of VDAC protein level SKBR-3 and NIH-3T3. Data are normalized to the β-actin and expressed as means ± SEM (n = 3) and analyzed with One-Way (*** $p < 0.001$). (C) VDAC1 mRNA expression evaluated by quantitative RT-PCR in tumor and paired non-tumor cells (*p<0,05).

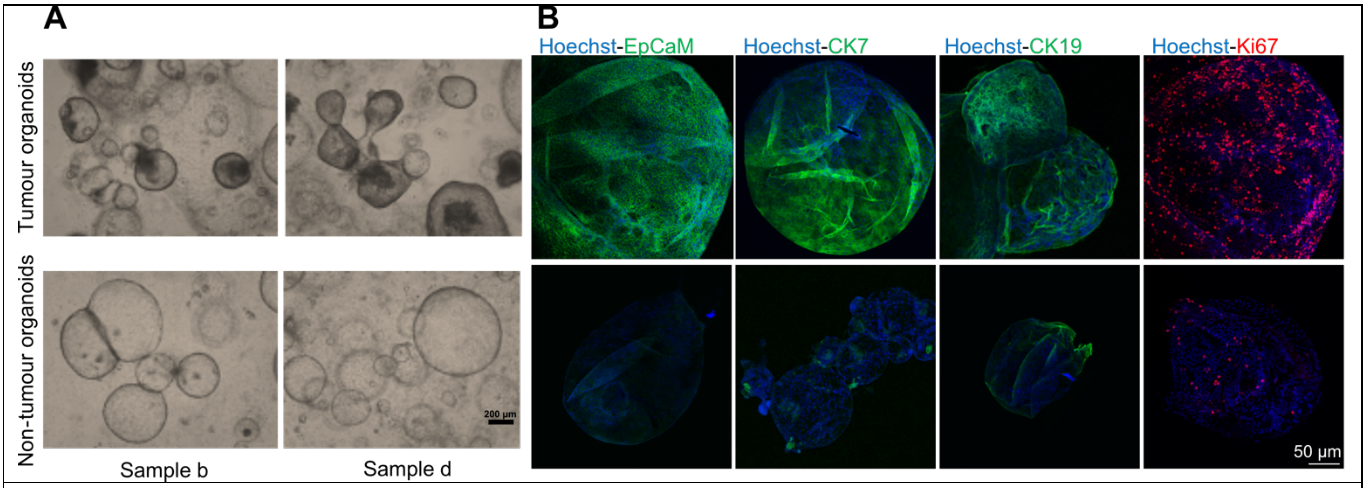


**Figure S4. Morphological and immunochemical validation of iCCA and paired non-tumor organoids**
A. Representative bright-field images of tumor and paired non-tumor organoids from two different iCCA patients. Scale bar: 200 μm. B. Immunofluorescence analysis for iCCA markers EpCaM (green), CK7 (green), and CK19 (green) and Ki67 (red) in confocal microscopy. Nuclei were counterstained with Hoechst33342 (blue). Scale bar: 50 μm

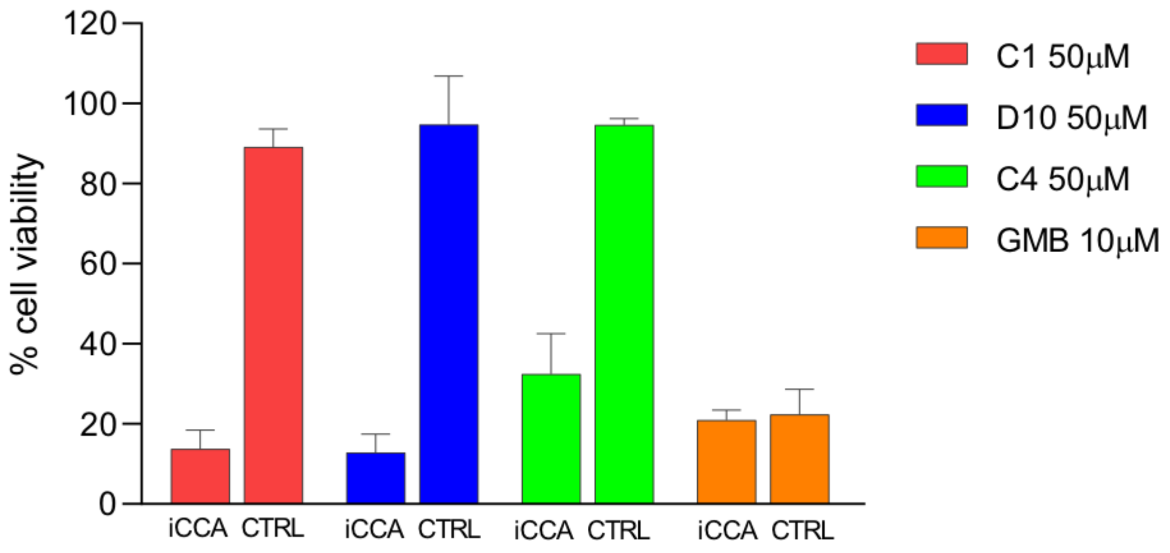


**Figure S5. Gemcitabine effect on iCCA organoids and non-tumor samples**
Gemcitabine (GMB) effect on iCCA organoids and non-tumor samples (CTRL) compared to VA molecules treatment effect (C1, D10, and C4). GMB exhibits pronounced effectiveness on both iCCA and control organoids, even at concentrations that are five-fold lower than those of VA molecules. This highlights the superior chemical potency of VA molecules in comparison to standard drugs.

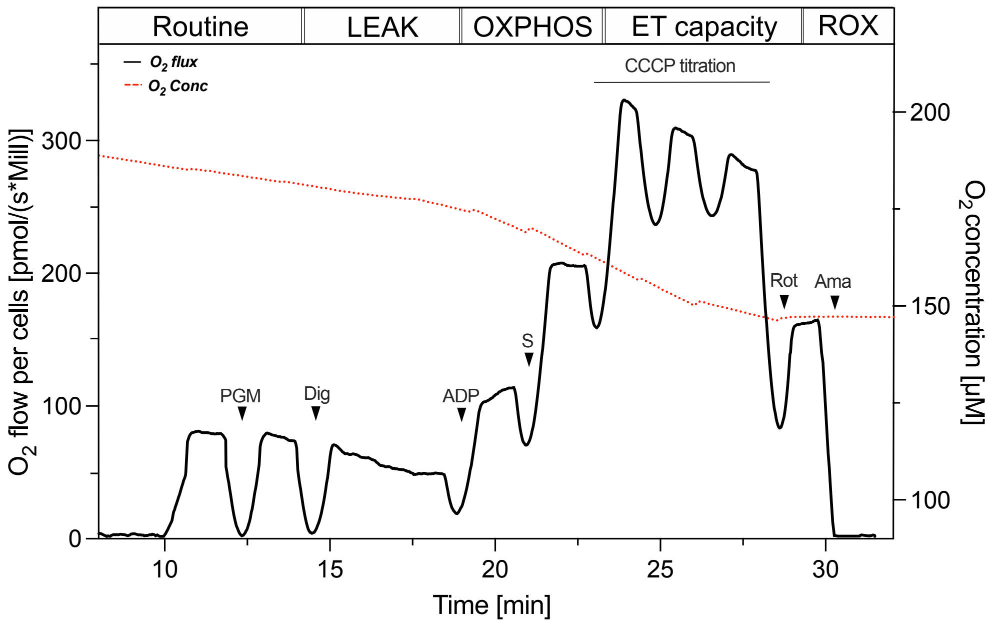


**Figure S6. Oxygen consumption in NIH - 3T3 cell line**. A representative curve of mitochondrial respiratory profile of NIH 3T3 untreated along with the SUIT protocol used in this work (P, pyruvate; M, malate; G, glutamate; Dig, digitonin; S, succinate; Rot, rotenone; Ama, antimycin).

| T(i) [°C] | | | | | | | | |
|---|---|---|---|---|---|---|---|---|
| | ***WT*** | ***N238K*** | ***N238F*** | ***N238R*** | ***R218Y*** | ***N207R*** | ***K236A*** | ***F18A*** |
| ***w/o DMF*** | 59.43 ± 0.03 | 59.74 ± 0.33 | 59.33 ± 0.31 | 59.37 ± 0.26 | 59.35 ± 0.04 | 59.39 ± 0.14 | 59.75 ± 0.19 | 59.74 ± 0.10 |
| ***0.5% DMF*** | 55.59 ± 0.61 | 55.85 ± 0.23 | 56.80 ± 0.35 | 55.50 ± 0.20 | 56.51 ± 0.50 | 56.44 ± 0.10 | 55.19 ± 0.14 | 55.87 ± 0.30 |

**Table S1**
The inflection temperature (T(i)). The table shows the T(i) values of the recombined mVDAC1 WT and mutants without DMF or pre-treated with 0.5% DMF. Data are reported as mean ± SEM of n=3 independent experiments.

| | ***SKBR-3*** | | ***NIH - 3T3*** | |
|---|---|---|---|---|
| | ***DMSO*** | ***D11*** | ***DMSO*** | ***D11*** |
| ***LEAK*** | 28.22 ± 9.52 | 14.12 ± 2.92 | 49.40 ± 2.21 | 48.98 ± 0.70 |
| ***N-Pathway*** | 79.46 ± 11.96 | 44.62 ± 12.13 | 117.36 ± 4.30 | 121.67 ± 5.84 |
| ***NS-Pathway*** | 144.36 ± 12.1 | 83.06 ± 10.18 | 237.52 ± 4.22 | 238.43 ± 4.34 |
| ***ET-Capacity*** | 242.1 ± 11.68 | 119.45 ± 5.40 | 299.61 ± 12.41 | 294.22 ± 7.06 |
| | ***DMSO*** | ***D10*** | ***DMSO*** | ***D10*** |
| ***LEAK*** | 23.70 ± 3.32 | 17.98 ± 2.22 | 52.06 ± 1.15 | 51.69 ± 1.33 |
| ***N-Pathway*** | 80.45 ± 3.67 | 47.39 ± 12.94 | 120.66 ± 5.33 | 119.88 ± 7.55 |
| ***NS-Pathway*** | 138.2 ± 15.86 | 101.01 ± 9.22 | 236.86 ± 13.41 | 241.69 ± 7.96 |
| ***ET-Capacity*** | 202.6 ± 15.82 | 142.03 ± 25.32 | 308.24 ± 8.25 | 306.91 ± 10.03 |
| | ***DMSO*** | ***C1*** | ***DMSO*** | ***C1*** |
| ***LEAK*** | 23.95 ± 5.13 | 24.52 ± 0.85 | 54.72 ± 1.85 | 55.24 ± 2.28 |
| ***N-Pathway*** | 87.50 ± 3.01 | 70.04 ± 7.82 | 122.64 ± 8.51 | 123.89 ± 9.99 |
| ***NS-Pathway*** | 142.2 ± 10.01 | 122.29 ± 6.37 | 250.40 ± 9.55 | 250.11 ± 17.47 |
| ***ET-Capacity*** | 254.64 ± 3.87 | 212.40 ± 11.75 | 300.22 ± 5.08 | 294.37 ± 6.18 |
| | ***DMSO*** | ***C4*** | ***DMSO*** | ***C4*** |
| ***LEAK*** | 30.99 ± 2.20 | 27.90 ± 6.91 | 44.55 ± 1.38 | 43.86 ± 0.51 |
| ***N-Pathway*** | 72.06 ± 7.46 | 58.91 ± 9.27 | 125.42 ± 1.77 | 123.58 ± 5.72 |
| ***NS-Pathway*** | 136.37 ± 2.77 | 111.36 ± 8.10 | 268.37 ± 3.68 | 263.64 ± 9.60 |
| ***ET-Capacity*** | 284.32 ± 3.34 | 233.48 ± 1.36 | 322.86 ± 7.07 | 329.28 ± 2.66 |
| | ***DMSO*** | ***C6*** | ***DMSO*** | ***C6*** |
| ***LEAK*** | 26.43 ± 4.24 | 22.95 ± 2.88 | 50.11 ± 1.90 | 48.95 ± 1.98 |
| ***N-Pathway*** | 78.69 ± 6.18 | 47.85 ± 7.33 | 119.38 ± 3.41 | 119.10 ± 3.39 |
| ***NS-Pathway*** | 141.43 ± 3.27 | 120.03 ± 6.64 | 258.16 ± 17.49 | 255.06 ± 14.68 |
| ***ET-Capacity*** | 275.52 ± 1.76 | 240.36 ± 7.33 | 308.19 ± 21.38 | 229.84 ± 21.30 |

**Table S2**. Oxygen flux calculated for each respiratory state corrected for the ROX respiration in untreated (DMSO) and VA-molecules treated cell lines (SKBR-3 and NIH-3T3). Data are reported as mean ± standard deviation of n = 3 independent experiments.

| Sample ID | Sex | Age (Years) | Liver Disease | Tumour stage | Differentiation grade | Tumour size (cm) | Vascular invasion |
|---|---|---|---|---|---|---|---|
| Sample 01 | male | 74 | NASH | T2 | G2 | >5 | yes |
| Sample 02 | female | 69 | none | T1 | G3 | >5 | no |
| Sample 03 | female | 82 | none | T1-T2 | G3 | <5 | no |
| Sample 04 | male | 76 | HCV | T1 | G2 | <5 | no |
| Sample 05 | male | 69 | none | T2 | G2 | <5 | yes |

**Table S3. iCCA patient clinical data.** Abbreviations: NASH: non-alcoholic steatohepatitis; HCV: hepatitis C virus

**Supplementary Methods**

*Immunofluorescence staining*

In order to perform an immunofluorescence characterization, organoids were fixed in cold 4% paraformaldehyde (Sigma-Aldrich) for 40 minutes at 4°C and subsequently resuspended in PBS with 0.1% Tween20 (Applichem) for 10 minutes at 4°C for the permeabilization phase. To reduce background non-specific staining, organoids were blocked in PBS with 0.1% Triton X-100 (Sigma-Aldrich) and 2% Bovine Serum Albumin (Pan Biotech) solution, for 1h at room temperature, and then primary antibodies Epithelial Cell Adhesion Molecule (EpCaM, 1:250; # 53-8326-42, Invitrogen), cytokeratine 19 (CK19, 1:250; #MA1-06329, Invitrogen), cytokeratine 7 (CK7, 1:250; #NBP2-44814, Novus Biologicals), and Ki67 (1:250; #MA5-14520, Invitrogen) were incubated overnight at 4°C. Fluorochrome-labeled secondary antibodies (1:500; #A-11001, and #A-21244, Invitrogen) in PBS solution with 0.1% Triton X-100 and 0.5% BSA were incubated for 1h at room temperature. Nuclei were counterstained with Hoechst33342 (#H1399, Invitrogen). Confocal images were captured on a Leica SP5 inverted confocal microscope (Leica).
